# Deletion induced splicing in *RIC3* drives nicotinic acetylcholine receptor regulation with implications for endoplasmic reticulum stress in human astrocytes

**DOI:** 10.1101/2022.07.18.500445

**Authors:** Navneesh Yadav, B. K. Thelma

**Author notes:** **Correspondence** B.K. Thelma, Professor, Department of Genetics, University of Delhi South Campus, Benito Juarez Marg, New Delhi – 110021, India.

## Abstract

Nicotinic acetylcholine receptor (nAChR) dysregulation in astrocytes is reported in neurodegenerative disorders. Modulation of nAChRs through agonists confers protection to astrocytes from stress but regulation of chaperones is unclear. Resistance to inhibitors of cholinesterase 3 (*RIC3*) is a potential chaperone of nAChRs but poorly studied in humans. We characterized *RIC3* in astrocytes derived from an isogenic wild-type and a Cas9 edited ‘del’ human iPSC line harboring a 25bp homozygous deletion in exon2. Altered *RIC3* transcript ratio due to deletion induced splicing and an unexpected gain of α7nAChR expression were observed in ‘del’ astrocytes. Transcriptome analysis showed higher expression of neurotransmitter/G-protein coupled receptors mediated by cAMP and calcium/calmodulin-dependent kinase signaling. Functional implications of these observations were examined using tunicamycin induced ER stress. Wild-type astrocyte stress model showed cell cycle arrest, *RIC3* upregulation, reduction in α7nAChR surface levels but increased α4nAChR surface expression. Conversely, tunicamycin treated ‘del’ astrocytes showed a comparatively higher α4nAChR surface expression and upsurged cAMP signaling. In addition, reduced expression of stress markers CHOP, phospho-PERK and lowered *XBP1* splicing in western blot and qPCR, validated by proteome-based pathway analysis indicated lowered disease severity. These findings indicate i) a complex RNA regulatory mechanism via exonic deletion induced splicing; ii) RIC-3 as a disordered protein having contrasting effects on co-expressed nAChR subtypes under basal/stress conditions; and iii) *RIC3* as a potential drug target against ER stress in astrocytes for nicotine related brain disorders. Furthermore, cellular rescue mechanism through deletion induced exon skipping possibly opens up ASO based therapies for tauopathies.

## 1. INTRODUCTION

Maturation of membrane proteins is a multi-step quality control process wherein a wide range of molecular chaperones assist in the folding of various ion channels, neurotransmitter receptors and G-protein coupled receptors (GPCRs) required for brain functioning ^1,2^. Among these, nicotinic acetylcholine receptors (nAChRs), the well studied membrane receptors are ligand-gated ion channels expressed in the brain where they control diverse physiological and behavioural functions such as neurotransmitter release, maintenance of synaptic transmission and plasticity ^3,4^. Functional upregulation of nAChRs in tobacco addiction has been documented early on ^5^; and their dysregulation contributing to the pathophysiology of various neurodegenerative and neuropsychiatric disorders including Alzheimer’s disease (AD), Parkinson’s disease (PD), Schizophrenia (SZ) etc. are also reported ^6–9^. Loss of antioxidant function, glutamate uptake etc. due to apoptosis; or a gain of neurotoxic function via neuroinflammation in astrocytes, the most abundant non-neuronal cell type in the brain, have been shown to result in disruption of neuronal function and neurodegeneration ^10,11^. This is expected considering astrocytes are involved in normal synapse formation, pruning and homeostasis ^12^, and also express functional nAChRs which indirectly regulate many neuronal events, such as neurotransmitter release and synaptic plasticity ^13^. Furthermore, increased α7 nAChRs in astrocytes for example, are associated with Aβ toxicity in AD ^14^. Conversely, α7 nAChR agonists PNU-282987 and nicotine, a pharmacological chaperone of nAChRs have been shown to confer protection to astrocytes from cellular stress by activating α7 nAChRs ^15,16^. These observations are interesting but regulation of cholinergic receptors in astrocytes is poorly understood. Elucidating these through modulation of their chaperones may be an effective approach.

*RIC3* (11p15.4) which codes for resistance to inhibitors of cholinesterase 3, and involved in folding and assembly of specific 5-hydroxytryptamine type 3 receptors and various nAChR subtypes, especially homomeric α7 receptors ^17,18^ is one such potential chaperone. Importance of *RIC3* has been amply demonstrated through genetic studies in *C*.*elegans* but knowledge of its function have relied on heterologous expression studies in different cell lines such as COS and HEK293 ^19^; posing some limitations. On the other hand, CRISPR/Cas9 genome editing in human iPSCs which enables generation of isogenic cellular models are powerful contemporary tools for characterization of such genes and also for drug discovery ^20^. In this study, we aimed at characterization of *RIC3* in astrocytes using wild-type and Cas9 edited gene and employing targeted and hypothesis free approaches. Of the several human iPSC lines with *RIC3* edits generated previously in the laboratory (manuscript under revision), a ‘del’ line with a 25bp homozygous deletion in exon 2 which is expected to result in gross structural and functional changes in RIC-3 was selected for this study. An unexpected altered *RIC3* transcript ratio due to 25bp deletion induced splicing; and gain of α7 nAChR surface expression with increased calcium/calmodulin-dependent kinase/cAMP signaling was evidenced in ‘del’ astrocytes. Efforts to explore the functional relevance, if any, of this observation under ER stress in ‘del’ astrocytes, showed increased surface expression of α4 nAChR subunit and cAMP signaling. Resultant lowered disease severity suggestive of *RIC3* as a drug target for nicotine related brain disorders emerged as a novel lead.

## 2. MATERIALS AND METHODS

### 2.1 Ethics statement

This work was carried out in accordance with the guidelines approved by the Institutional Committee for Stem Cell Research, University of Delhi South Campus (ICSCR-UDSC#2018/01).

Details of the cell lines/cell types used; and methodology for *RIC3* characterization are detailed below.

### 2.2 Maintenance of cell lines

A well characterized CRISPR/Cas9 edited ‘del’ iPSC line carrying a homozygous 25bp deletion in exon 2 of *RIC3* along with the isogenic unedited wild-type iPSC line (Gibco #A18945) were used in this study. The ‘del’ line was generated previously in the laboratory by utilizing the microhomology mediated end joining dsDNA repair mechanism induced by CRISPR/Cas9 (manuscript under revision). The ‘del’ line had a normal karyotype (46, XX) without any Cas9 mediated off-targets as evaluated by Sanger sequencing of predicted off-targets, array CGH and exome sequencing; and expressed commonly tested pluripotency markers (data not provided). In the present study, iPSCs were cultured in six well plates on geltrex matrix in 2ml of stemflex medium at 37°C and 5% CO_2_. Medium was changed every alternate day and passaged (1:6) as small clumps every 3 to 4 days using versene. Mycoplasma contamination in the running cultures was screened routinely by PCR. For confirmation of 25bp deletion in *RIC3* in ‘del’ line prior to astrocyte differentiation, genomic DNA was isolated using DNA purification kit (Promega #A1120) following manufacturer’s instructions. PCR was carried out using Q5 High-Fidelity DNA Polymerase (NEB #M0491S) with primers (Supplementary Table 1) and Sanger sequenced using reverse primer at the Central Instrumentation Facility (CIF, UDSC). All the reagents were obtained from ThermoFisher Scientific, US unless stated otherwise and primers were from Merck, India.

### 2.3 Astrocyte differentiation from iPSC derived neural stem cells and characterization

#### a) Neural induction of iPSCs

In both ‘del’ and wild-type iPSC cultures with ∼15-20% confluence, StemFlex medium was replaced with PSC Induction Medium containing Neurobasal medium and Neural Induction Supplement. For neural induction, medium was changed every alternate day and daily from day 5 when cells were ∼90% confluent. The growing primitive Neural Stem Cells (NSCs) were then dissociated using accutase and passaged on geltrex coated plates in NSC expansion medium containing 50% Neurobasal medium, 50% advanced DMEM/F12 and Neural Induction Supplement. Confluent NSCs were passaged (1:3) every 4 to 5 days using accutase and characterized as detailed below.

#### b) RNA isolation and semi-quantitative real time PCR (RT-PCR)

For characterization of NSCs generated from both lines, total cellular RNA using ∼1×10^6^ cells from one well of a six-well plate was isolated using TRIzol reagent followed by Direct-zol RNA MiniPrep Plus kit purification (Zymo Research # R2070). RNA was quantified using Qubit RNA HS Assay kit and cDNA synthesised from ∼2µg of RNA using SuperScript IV VILO Master Mix kit following gDNA removal with ezDNase. Expression of neural markers *NESTIN* and *SOX1* in NSCs was tested by RT-PCR using primer pairs (Supplementary Table 1) designed from CDS region and SYBR green reagent, as per manufacturer’s instructions on a QuantStudio 6 system (Applied Biosystems). The raw data obtained from two independent experiments, each done in triplicates were analysed and the relative gene expression normalised with *18S* was plotted (ΔCt method).

#### c) Immunocytochemistry and imaging

For immunostaining of NSCs, cells were grown in geltrex coated 8 chamber slides and fixed using 4% PFA in DPBS for 15 mins, permeabilized using 1% Saponin in DPBS for 15 mins, blocked with 3% BSA for 30 mins at RT and incubated overnight at 4°C with primary antibodies (NESTIN and SOX-2). Cells were washed with DBPS (3X) and incubated with the respective secondary antibodies (Supplementary Table 2) for two hours at RT and DAPI stained. Mounted slides were scanned under Leica SP5 Confocal Laser Scanning Microscope at CIF, UDSC and analysed using Leica software; and Adobe Photoshop CS5 was used for image processing.

#### d) Astrocyte differentiation and characterization

For astrocyte differentiation, well characterized NSCs at passage six were dissociated using accutase and plated onto geltrex-coated six well plates at a density of 5×10^4^ cells per cm^2^ in an astrocyte differentiation medium consisting of DMEM+GlutaMAX, N-2 Supplement and Fetal Bovine Serum. Confluent cultures were passaged (1:3) until day 30 and later stained with GFAP (Supplementary Table 2), an astrocyte marker.

### 2.4 *RIC3* characterization under basal condition in wild-type and ‘del’ astrocytes

#### a) qPCR

*RIC3* qPCR was performed with cDNA from astrocytes from the respective cell lines using Q5 High-Fidelity DNA Polymerase with primers designed from exons 1, 3 and 6 (Supplementary Table 1). Amplicons were checked on 2.5% agarose gel, eluted using GeneJET Gel Extraction kit and confirmed by Sanger sequencing.

#### b) Protein isolation, quantification and western blot

Cytosolic and membrane proteins from the respective lines were isolated using Mem-PER Plus Membrane Protein Extraction Kit in presence of Halt Protease and phosphatase inhibitor Cocktail as per manufacturer’s instructions. Total protein concentration was estimated using Pierce BCA Protein Assay kit. For western blot, equal amounts of protein were loaded onto 10% SDS PAGE followed by 3 hours fast transfer onto PVDF membrane. Membrane was blocked overnight using 5% BSA at 4°C and subsequently incubated overnight with primary antibody against CHRNA7 (Neuronal acetylcholine receptor subunit alpha-7) with GAPDH (Glyceraldehyde-3-phosphate dehydrogenase) as internal control. Membrane was washed with 1X TBST and incubated with respective HRP conjugated secondary antibody for two hours and washed before developing the blots using chemilumiscence kit (Thermo Fisher Scientific, US). Details of all the primary and secondary antibodies used for western blot is provided in Supplementary Table 2.

### 2.5 *RIC3* characterization under stress in wild-type and ‘del’ astrocytes

Astrocyte model of disease was developed using tunicamycin. Mature astrocytes at day 30 were treated with tunicamycin (5µg/ml; Sigma #SML1287) for 24-48 hours for stress induction. Wild-type and ‘del’ astrocytes with and without tunicamycin treatment in triplicates were grown in 6 well plates and used for transcriptomics, western blot and proteomics as detailed below.

#### a) Transcriptomics

For RNA sequencing, the purity/integrity of isolated RNA was checked on UV/VIS spectrophotometer (Qiagen, US) and RNA HS ScreenTapes system (Agilent Technologies, US). For library preparation, rRNA was removed using RiboCop rRNA depletion kit (Lexogen, US); cDNA synthesized from enriched RNA using reverse transcriptase and random primers; converted into double-stranded DNA and enriched with PCR. PCR products were then purified and checked for fragment size distribution using D1000 DNA ScreenTapes (Agilent Technologies, US). Sample libraries were multiplexed and sequenced on a NovaSeq 6000 (Illumina, San Diego, California, US) using paired-end sequencing (150bp) at a commercial facility (MedGenome Labs, Bengaluru, India). On an average, >40 million trimmomatic filtered reads were generated and aligned to the human genome (hg19) using splice aware aligner, HISAT2 ^21^. The total number of uniquely mapped reads were counted using feature counts and subjected to differential gene expression (DGE) using DeSeq2 ^22^. Differentially expressed genes (p-value≤0.05;±1 fold change) were considered as significant and taken forward for analysis.

#### b) Western blot and Proteomics

Expression of CHRNA7, CHRNA4 (Neuronal acetylcholine receptor subunit alpha-4), CHOP (C/EBP-homologous protein), phospho-PERK (Protein kinase RNA-like ER kinase) and PIK3R3 (Phosphoinositide-3-kinase regulatory subunit 3) with GAPDH as internal control were checked with antibodies on western blots using protocol described above (4b). For proteomics, cells were resuspended in pre-heated (70°C) 6M Guanidine Hydrochloride buffer containing 1X Halt Protease and phosphatase inhibitor Cocktail; vortexed for 2 minutes at RT; boiled at 100°C for 3 minutes; centrifuged at 15,000 rpm for 10 mins at RT; and supernatant containing proteins quantified using Pierce BCA Protein Assay kit. The protein samples were run on Q-Exactive Plus Orbitrap Mass Spectrometer at a commercial facility (VProteomics, New Delhi, India). RAW files were analysed with Proteome Discoverer (v2.2, Thermo Fisher Scientific, USA) against the Uniprot *homo sapiens* database using SEQUEST and AMANDA. Label-free quantification was done using Proteome Discoverer software and statistics performed using Perseus (v1.6.2.3) software. Only proteins which were differentially expressed (adjusted p-value≤0.05;±1 fold change) were considered for analysis.

#### c) Data analysis

GO enrichment analysis and transcription factor identification were done for the set of significant differentially expressed genes from transcriptome data (2.5a) and proteins (2.5b) obtained using DAVID ^23^. The Cytoscape plugin Enrichment map and GOPLOT R package was used to visualise top BP (biological process), CC (cellular component), MF (molecular function), Reactome and Kyoto Encyclopedia of Genes and Genomes (KEGG) terms (adjusted p-value≤0.05) ^24^.

#### d) Validation by RT-PCR, qPCR and immunostaining

Differentially expressed functionally relevant genes were validated by SYBR green based RT-PCR. Data were analysed, relative gene expression calculated with *GAPDH* as endogenous control and shown as fold change (2^-^ΔΔCt method). *RIC3* expression was validated by qPCR using Q5 High-Fidelity DNA Polymerase with primers designed from exons 1 and 4; and immunostaining using RIC-3 primary antibody. For *XBP1*, a stress gene, qPCR was performed using Q5 High-Fidelity DNA Polymerase and splicing, if any, was confirmed by checking the PCR products on a 2% agarose gel. All primers and antibodies used are listed in Supplementary Tables 1 and 2 respectively.

## 3. RESULTS

### 3.1 Efficient differentiation of iPSC derived NSCs to astrocytes

High quality NSCs were generated from an isogenic wild-type iPSC line and a ‘del’ iPSC line having a homozygous 25bp deletion in exon 2 of *RIC3* (Fig. 1a) using serum-free neural induction and expansion media. Compact NSCs with uniform morphology (Fig. 1b) expressed neural markers *NESTIN* and *SOX1*, confirmed by SYBR green based RT-PCR (Fig. 1c); and stained positive for SOX2 and NESTIN markers (Fig. 1d). NSCs (at passage 6) were differentiated into astrocytes and confirmed by GFAP staining (Fig. 1e). Morphological feature of astrocytes with characteristic star shaped structure was visualized at passages 2-3; and between passages 4-5, these manifested mainly flattened polygonal to fusiform morphology (Supplementary Fig. 1). Of note, no phenotypic differences were seen between wild-type and ‘del’ astrocytes, indicating that deletion in *RIC3* did not affect the differentiation process.

**Fig. 1.**
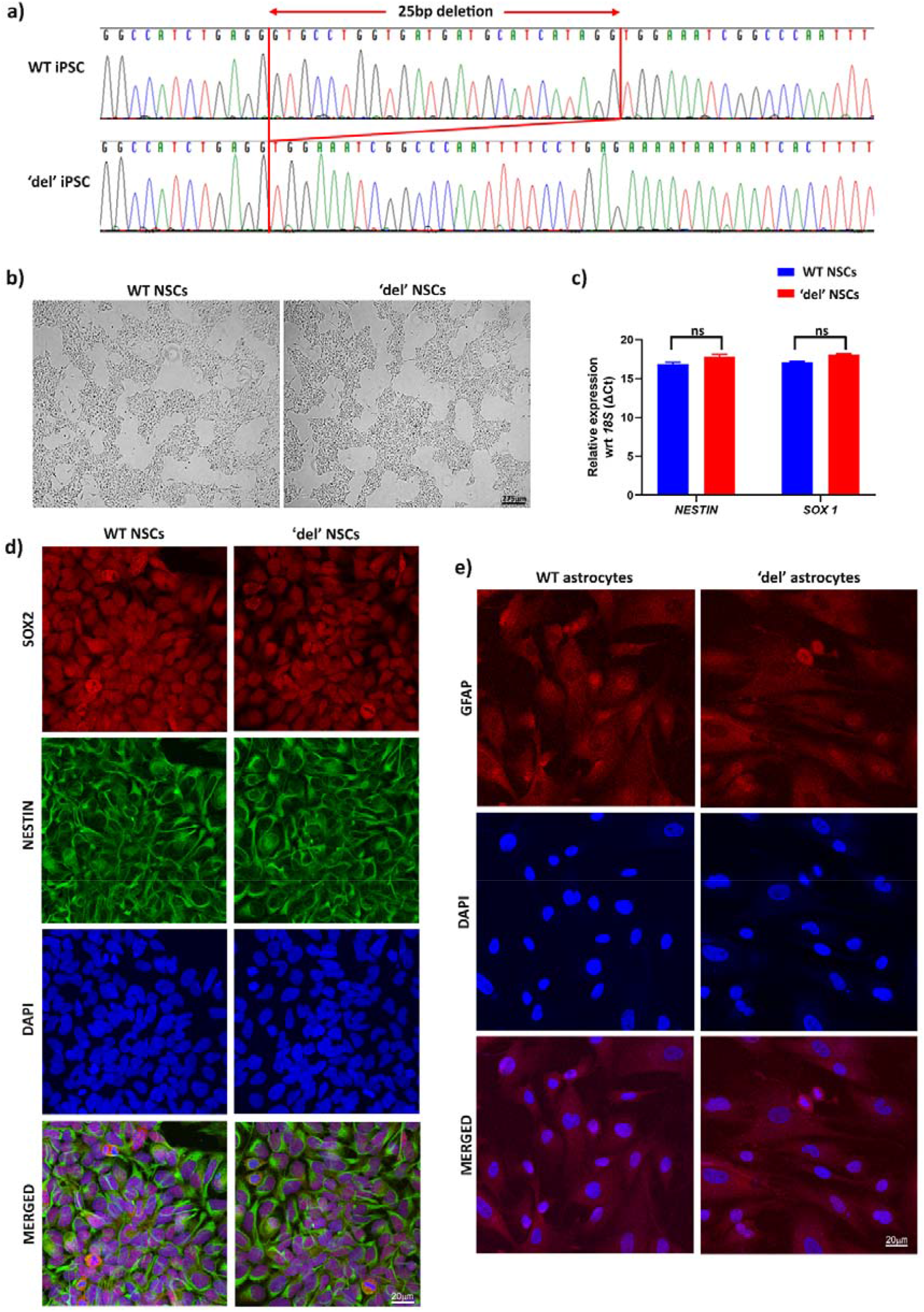
Generation and characterization of wild-type (WT) and ‘del’ iPSC derived neural stem cells (NSCs) and astrocytes **a)** Electropherograms of Sanger sequencing showing a part of *RIC3* exon 2 in WT iPSC line along with 25bp deletion (region denoted with red lines) in ‘del’ iPSC line obtained using reverse primer; **b)** representative brightfield images of NSCs generated from WT and ‘del’ iPSCs showing uniform compact colony morphology at passage 3 (scale bar: 275μm); **c)** relative expression of neuronal markers *NESTIN* and *SOX1* normalized with *18S* (ΔCt method) in WT and ‘del’ NSCs; **d, e)** representative immunofluorescence confocal images (Leica SP5) showing **d)** WT and ‘del’ NSCs expressing Alexa Fluor 594 labelled SOX2; Alexa Fluor 488 labelled NESTIN; counterstained with DAPI; and merged (scale bar: 20μm); and **e)** WT and ‘del’ astrocytes stained with Alexa Fluor 594 labelled GFAP; DAPI; and merged (scale bar: 20μm). Note no phenotypic/molecular differences were seen between WT and ‘del’ NSCs and astrocytes.

### 3.2 *RIC3* characterization in astrocytes

In view of the role of *RIC3* in assembly and surface trafficking of nAChRs, functional characterization using gene specific and hypothesis free approaches were performed in wild-type and ‘del’ astrocytes and results are detailed below.

#### a) Splicing alterations due to 25bp deletion in RIC3 and increased α7 nAChR surface expression seen in ‘del’ astrocytes

In the wild-type astrocytes, two alternatively spliced annotated *RIC3* transcripts (Supplementary Fig. 2a) were observed in qPCR performed using primers from exons 1 and 3. However, in the ‘del’ astrocytes, using the same primers spanning the deletion site, a mixture of novel transcripts with i) 25bp deletion and ii) exon 2 skipping were observed (Fig. 2a); and confirmed by Sanger sequencing (Fig. 2b). A similar splicing alteration in ‘del’ astrocytes was observed when primers from exons 1 and 6 were used in qPCR essentially to check the presence and abundance of different *RIC3* transcripts (Fig. 2a). In addition, interestingly, an increased expression of an annotated shorter *RIC3* transcript and a novel transcript with exons 1, 5 and 6 were observed (Fig. 2a and Supplementary Figs. 2a, b). These findings of altered splicing were supported by *in-silico* analysis using ESEfinder v3.0 (http://krainer01.cshl.edu/cgi-bin/tools/ESE3/esefinder) ^25^ and EX-SKIP (https://ex-skip.img.cas.cz/) ^26^, which predicted a loss of multiple exonic splicing enhancer (ESE) binding sites in exon 2 with a higher chance of exon skipping in ‘del’ astrocytes (Supplementary Fig. 2c). RIC-3 is a known molecular chaperone of nAChRs and therefore, the effect of splicing and altered *RIC3* transcript ratio on the levels of homomeric α7 nAChR on surface was checked. Of note, a significant increase in the CHRNA7 levels was witnessed in membrane fractions isolated from ‘del’ as compared to wild-type astrocytes on western blot (Fig. 2c). This suggests that the 25bp deletion induced splicing due to loss of ESE binding sites and/or altered *RIC3* transcript ratio may have resulted in increased surface expression of α7 nAChR.

**Fig. 2.**
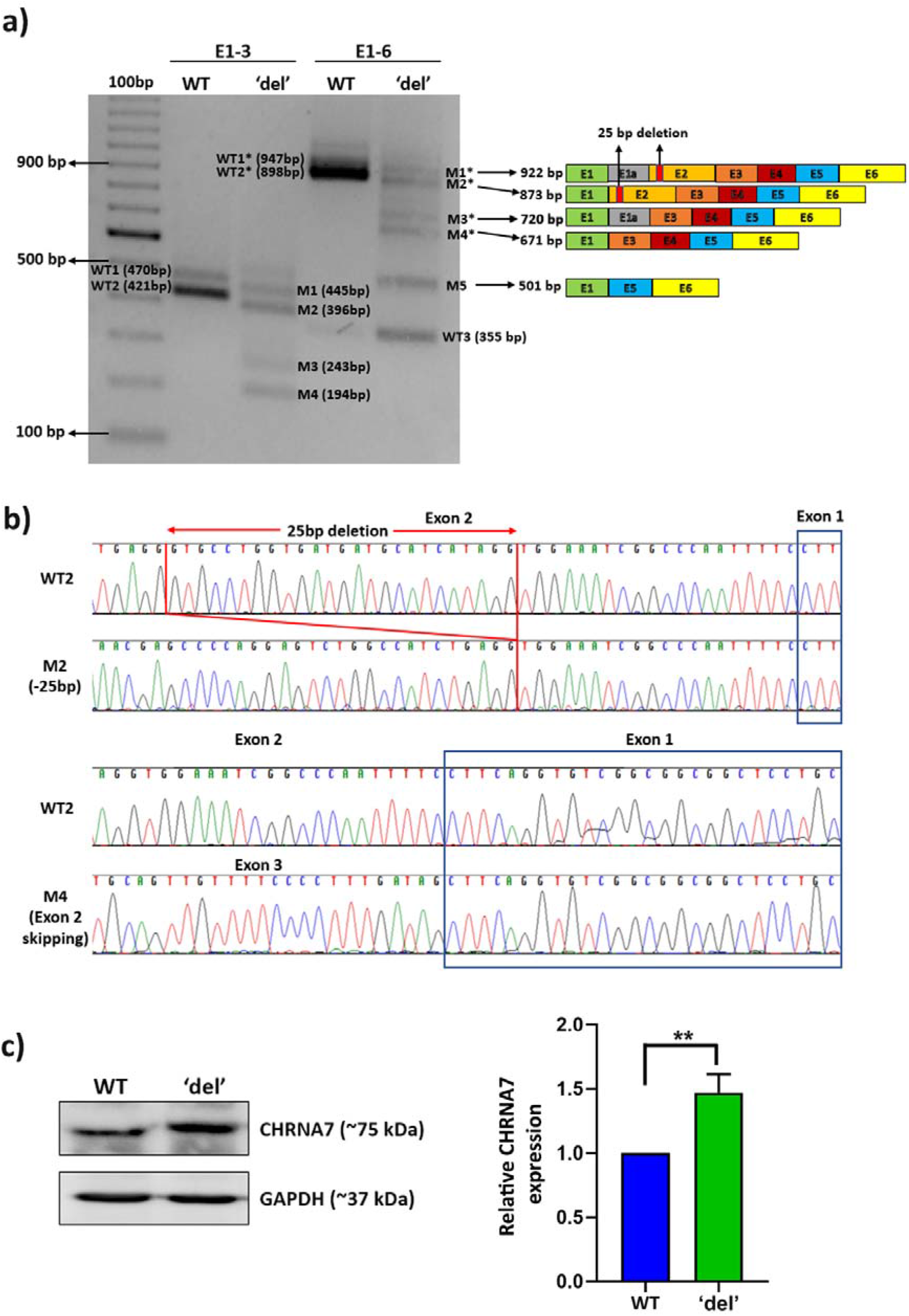
Characterization of *RIC3* in wild-type (WT) and ‘del’ astrocytes under basal condition **a)** qPCR profile and schematic presentation showing *RIC3* transcripts in WT and ‘del’ astrocytes captured using primers from exons 1-3 and 1-6. E1-3 shows two annotated *RIC3* transcripts WT1, 470bp; and highly expressed WT2, 421bp in WT astrocytes; and same transcripts but shorter due to 25bp deletion in exon 2, M1, 445bp; and M2, 396bp in ‘del’ astrocytes. Also seen are transcripts M3, 243bp; and M4, 194bp with exon 2 skipping due to mis-splicing. E1-6 shows three different full length annotated *RIC3* transcripts WT1*, 947bp; highly expressed WT2*, 898bp; and WT3, 355bp in WT astrocytes; and 25bp deletion mediated leaky splicing generating full length novel transcripts M1*, 922bp; M2*, 873bp; M3*, 720bp; and M4*, 671bp. Also seen is a novel transcript with total skipping of exons 2-4, M5, 501bp; and the highly expressed shorter annotated transcript WT3, 355bp in ‘del’ astrocytes; **b)** Sanger sequencing confirmation of the gel eluted fragments corresponding to WT2 and novel transcripts, M2 and M4 seen at (a) above; and **c)** Western blot showing expression of membrane bound CHRNA7 relative to GAPDH, internal control in WT and ‘del’ astrocytes. Note significantly enhanced expression of CHRNA7 in ‘del’ astrocytes compared to WT astrocytes observed in two independent blots (unpaired t-test, p-value<0.01).

#### b) Higher expression of ion-channels, neurotransmitter and GPCRs in ‘del’ astrocytes

To gain insight into the molecular changes associated with an increased expression of surface α7 nAChR in ‘del’ astrocytes, a hypothesis free approach of transcriptome analysis was adopted. Quality data having >90% reads aligned to genome hg19 with average Q30 ∼92% were analysed. DGE was performed using DeSeq2 which identified 773 upregulated and 682 downregulated genes in ‘del’ astrocytes as compared to wild-type astrocytes (p-value≤0.05;±1 fold change) (Fig. 3a); of which 61% (n=888/1455) were protein coding transcripts. Enrichment analysis based on GO, KEGG and Reactome terms identified using DAVID (adjusted p-value≤0.05) demonstrated significant perturbations in genes associated with a wide range of cellular functions including extracellular matrix organization, cell adhesion, signal transduction, cytokine-mediated signaling pathway, chemical synaptic transmission, regulation of NMDA receptor activity etc. (Supplementary Fig. 3a). Enrichment analysis of only the upregulated protein coding genes (n=442) revealed genes present in GO terms “plasma membrane”, “signal transduction”, “chemical synaptic transmission” and REACTOME terms “Neural system”, “Signaling by GPCR”, “Neurotransmitter receptors and postsynaptic signal transmission” and others (adjusted p-value≤0.05) (Fig. 3b). Specifically, an increased expression of genes coding for ion channels which included voltage-gated calcium, potassium, sodium/calcium, chloride channels; neurotransmitter receptors – glutamate, GABA, cholinergic muscarinic/nicotinic; and several G-protein coupled receptors (GPCRs) was evidenced. Furthermore, a higher expression of genes involved in calcium mediated exocytosis and release of neurotransmitters/cytokines was also noted. Among those involved in synaptic plasticity, an increased expression of calcium/calmodulin-dependent protein kinase genes, those involved in cAMP regulation and ERK/MAPK signaling pathway were seen. Upregulation of genes encoding cytokines and their receptors was noteworthy. UCSC_TFBS feature in DAVID also identified several transcription factors involved in calcium/calmodulin-dependent kinase signaling, cAMP signaling, inflammatory responses and entry into cell cycle (Fig. 3c; Supplementary Fig. 3b). Representative genes from the above categories are shown as Heat map (Fig. 3d) with their log2FC values mentioned elsewhere (Supplementary Table 3). A few among these, namely *CHRNA1, GABRQ, GRIN2A, GRIA1, CHRM2, GABBR2, GRIA3* (neurotransmitter receptors), *IL1B, IL6R* (cytokines/receptors), *MAPK13* and *SNCA* (synaptic plasticity) were validated by RT-PCR using cDNA obtained from an independent passage of wild-type and ‘del’ astrocytes (Fig. 3e). This hypothesis free analysis indicates a gain of calcium/calmodulin-dependent kinase and cAMP signaling in ‘del’ astrocytes through an increase in gene expression of various ion-channels, neurotransmitter receptors, GPCRs and genes involved in synaptic plasticity.

**Fig. 3.**
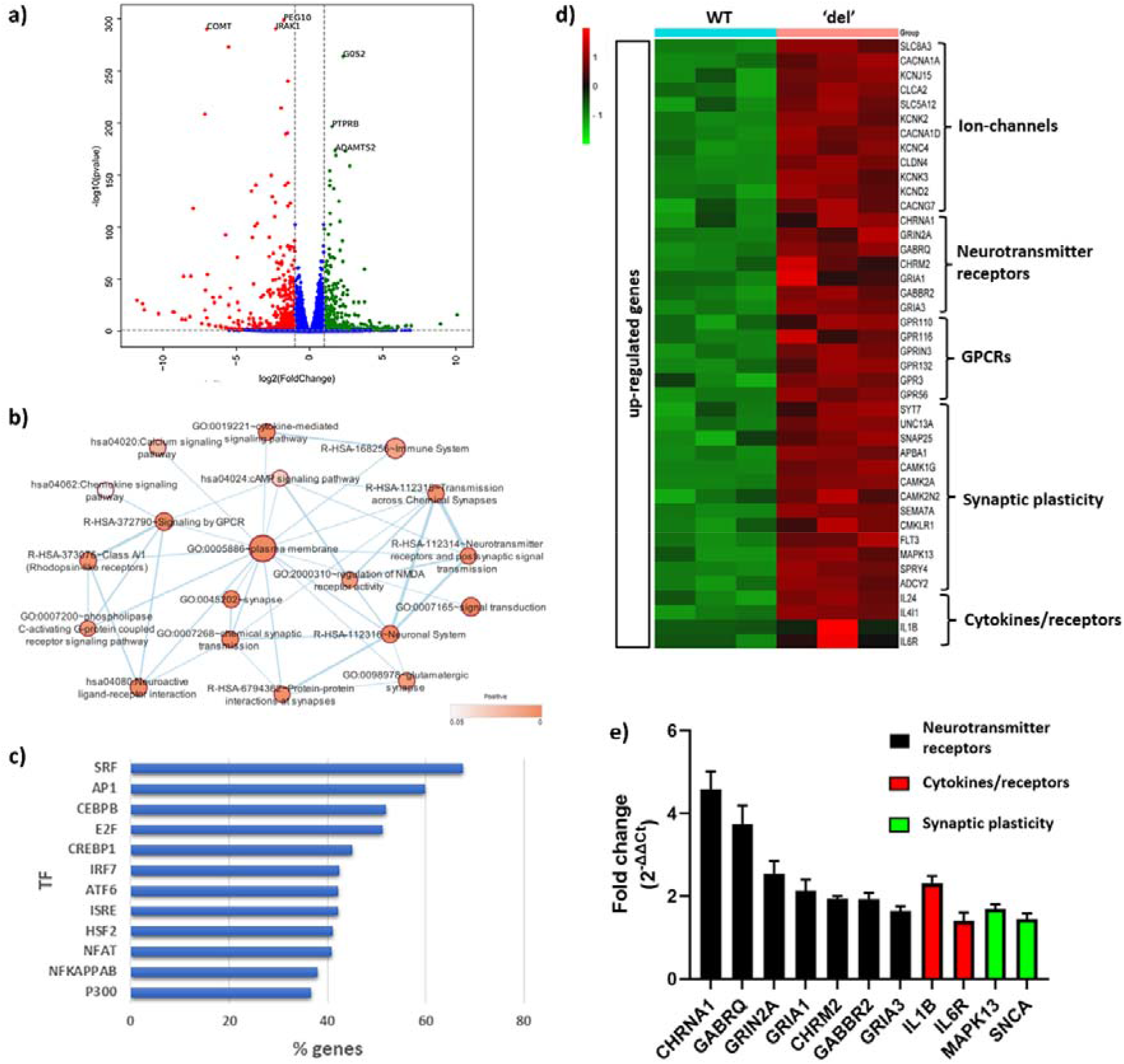
Comparative transcriptome analysis between wild-type (WT) and ‘del’ astrocytes under basal condition **a)** Volcano plot showing up (green, n=773) and down (red, n=682) regulated genes (each of the dots, p-value≤0.05;±fold change) in ‘del’ astrocytes compared to WT; **b)** enrichment analysis of significantly upregulated protein coding genes (n=442, p-value≤0.05;+1fold change) in ‘del’ astrocytes showing genes present in highly significant GO terms “plasma membrane”, “signal transduction”, “chemical synaptic transmission” and REACTOME terms “Neural system”, “Signaling by GPCR”, “Neurotransmitter receptors and postsynaptic signal transmission” and others (adjusted p-value≤0.05) identified using DAVID functional annotation tool. The color of the node and edge size correspond to the significance of the gene-set and number of genes overlapping between two connected gene-sets respectively; **c)** bar plot showing percentage of upregulated protein coding genes regulated by transcription factors involved in calcium/calmodulin-dependent kinase signaling (SRF), cAMP signaling (AP1, CREBP1, ATF6), inflammatory responses (CEPB, IRF7, ISRE, NFKAPPB) and entry into cell cycle (E2F) identified using UCSC_TFBS feature in DAVID; **d)** heat map constructed using selected genes observed in GO terms at (b) above; and **e)** validation of selected genes from candidate gene-sets at (b) above by RT-PCR normalized with *GAPDH*.

### 3.3 Functional implications of increased expression of surface receptor genes

Increased signaling likely via α7 nAChRs (preceding section 3.2b) in ‘del’ astrocytes was an unexpected finding. This novel observation which may have functional implications for nicotine related disorders was examined using tunicamycin induced ER stress, a commonly proposed pathomechanism in various neurodegenerative diseases. Such a disease model was established in wild-type astrocytes and findings were validated in tunicamycin treated ‘del’ astrocytes.

#### a) ER stress results in cell cycle arrest, increased unfolded protein response and higher RIC3 expression in tunicamycin treated wild-type astrocytes (disease state)

Tunicamycin treatment of wild-type astrocytes for 24 hrs. caused disruption in stellate shape of the astrocytes, visualized by brightfield microscopy (Fig. 4a); and cell death on continued treatment upto 48 hrs. (data not shown). Comparative transcriptome analysis of the tunicamycin treated (24 hrs.) and untreated wild-type astrocytes (in triplicates) revealed a total of 1918 upregulated and 1783 downregulated genes (p-value≤0.05;±1 fold change) (Fig. 4b); of which only 64.9% (n=2403) were protein coding transcripts. Enrichment analysis of these genes using DAVID identified an increase in expression of genes present in GO terms “endoplasmic reticulum unfolded protein response” and “IRE1-mediated unfolded protein response” (Fig. 4c and Supplementary Fig. 4), encompassing genes namely *HSPA5, DDIT3, DNAJC3, DNAJB9,11* and *EDEM1* etc. Higher expression of *CANX, CALR* and others involved in calnexin/calreticulin cycle, suggestive of altered calcium signaling was evident. On the other hand, a decrease in expression of genes involved in GO terms, “cell division”, “mitotic spindle organization” and “DNA replication” was observed (Fig. 4c and Supplementary Fig. 4). These included notable cell cycle genes namely *CDC20, CDK1, CDCA5, 8, MCM7, 10, BRCA1* and centromeric/mitotic spindle checkpoint genes, *CENPA, CENPE, CENPF, CENPL, CENPM, CENPN, CENPO, CENPU* among others. Selected genes representative of the above categories are shown as Heat map (Fig. 4d) with their log2FC values mentioned elsewhere (Supplementary Table 4). Interestingly, increased expression of *RIC3* was seen as confirmed by both RT-PCR and immunostaining which also showed RIC-3 aggregation (Figs. 4e and f). Similarly, RT-PCR results of unfolded protein response genes (*HSPA5, HSP40, CHOP*); calcium binding chaperones (*CALR, CANX*); cell cycle genes (*MCM7, CDC20, CENPA*); and *SNCA*, a known PD gene validated the transcriptome findings (Fig. 4e).

**Fig. 4.**
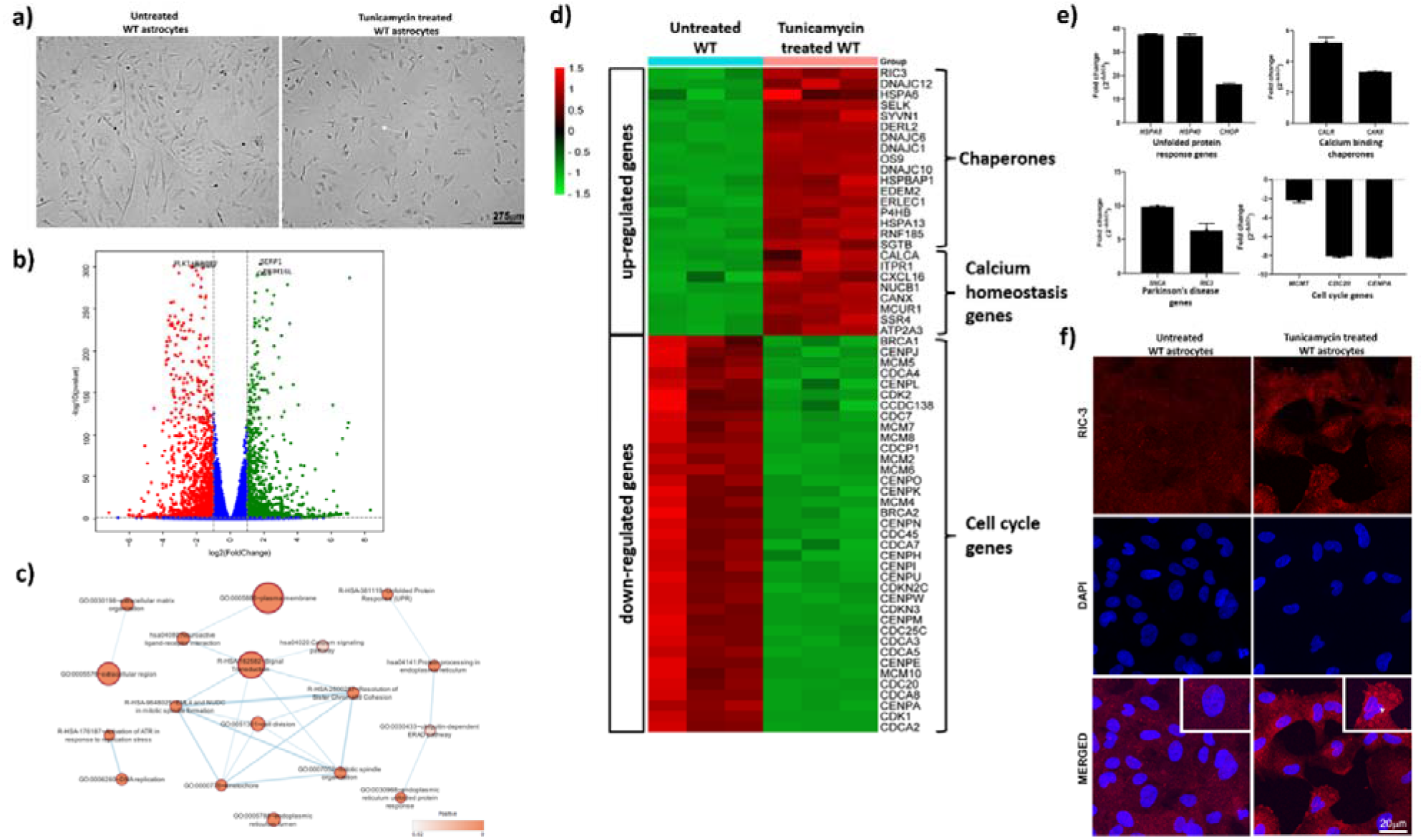
Establishment of tunicamycin induced ER stress astrocyte model (disease proxy) **a)** Representative brightfield images of wild-type (WT) astrocytes showing uniform flattened polygonal to fusiform morphology; and loss of projections (↑) due to tunicamycin treatment (24hrs) (scale bar: 275μm); **b)** volcano plot showing up (green, n=1918) and down (red, n=1783) regulated genes (each of the dots, p-value≤0.05;±fold change) in tunicamycin treated WT astrocytes compared to untreated WT; **c)** enrichment analysis of differentially expressed genes (p-value≤0.05;±1fold change) in tunicamycin treated WT astrocytes showing genes present in highly significant GO terms “plasma membrane”, “extracellular region”, “cell division” and REACTOME terms “signal transduction”, “unfolded protein response”, “resolution of sister chromatid cohesion” and others (adjusted p-value≤0.05) identified using DAVID functional annotation tool; **d)** heat map constructed using selected genes observed in GO terms at (c) above; **e)** validation of selected genes from candidate gene-sets at (c) above by RT-PCR normalized with *GAPDH*; and **f)** representative immunofluorescence confocal images (Leica SP5) showing increased Alexa Fluor 594 labelled RIC-3 expression and aggregation (arrow head); DAPI stained; and merged (scale bar: 20μm) in tunicamycin treated WT astrocytes.

#### b) Increased expression of surface α4 nAChR subunits and/or neurotransmitter/GPCRs in tunicamycin treated ‘del’ astrocytes reduces disease severity

Having established a cellular disease model in wild-type astrocytes as described in the preceding paragraph, the effect of altered *RIC3* transcript ratio on nAChRs regulation and expression of surface receptor genes were checked in tunicamycin treated ‘del’ astrocytes. Interestingly, an increase in *RIC3* mRNA levels (Fig. 5a), but a reduction in surface expression of α7 nAChR estimated by western blot in both tunicamycin treated wild-type and ‘del’ astrocytes as compared to untreated were observed (Fig. 5b). Conversely, a very low expression of CHRNA4, another nAChR subunit was seen in both untreated wild-type and ‘del’ astrocytes but increased with tunicamycin treatment. Interestingly, its expression was notably higher in tunicamycin treated ‘del’ astrocytes (Fig. 5b). On the other hand, PIK3R3, a signaling protein known to function downstream of α7 nAChR remained unchanged (Fig. 5b).

**Fig. 5.**
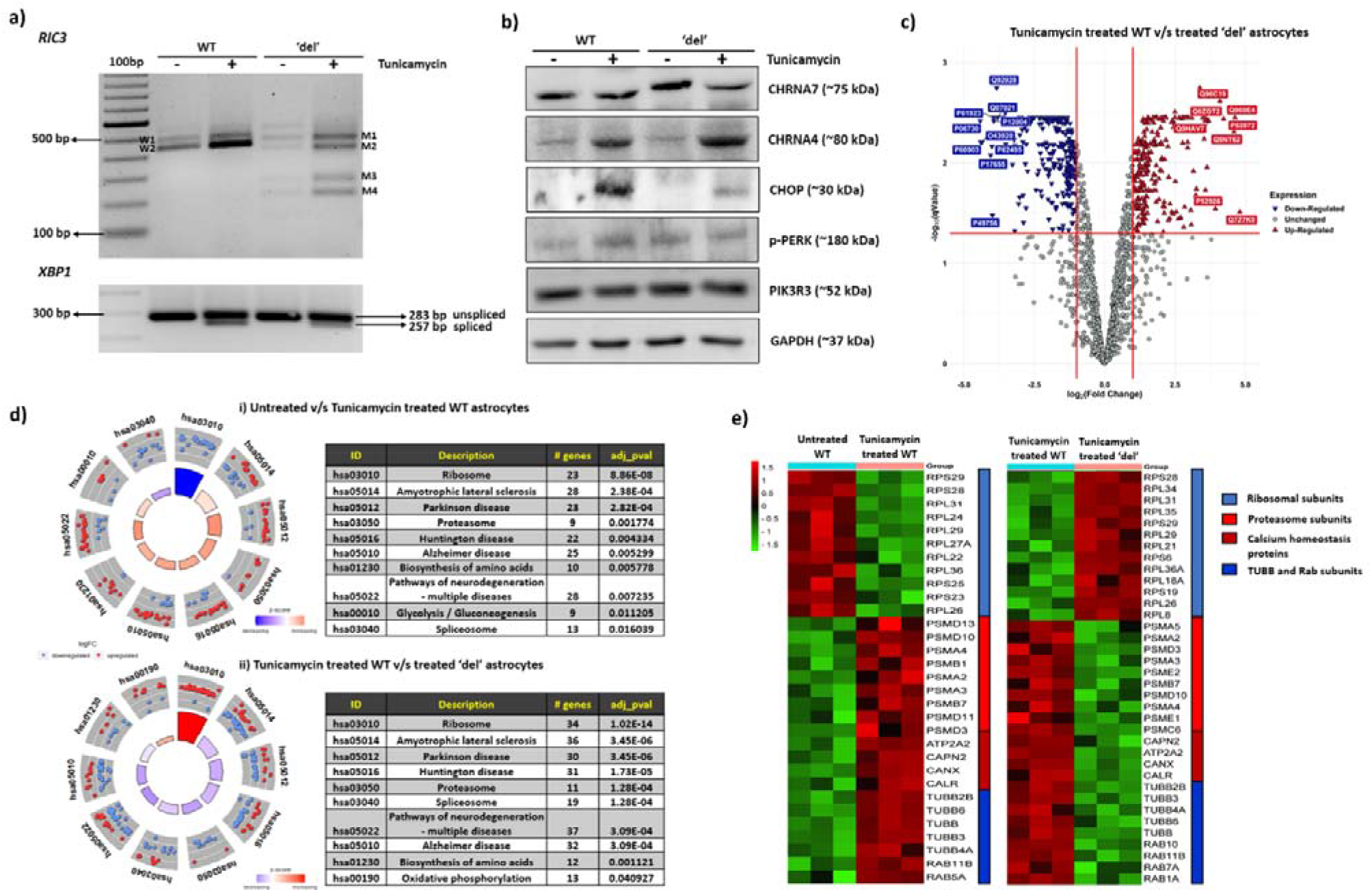
Characterization of *RIC3* in wild-type (WT) and ‘del’ astrocytes under tunicamycin treated conditions using gene based and proteomic approaches **a)** qPCR profile showing an overall increase in *RIC3* expression using primers from exons 1-4; and *XBP1* splicing in both tunicamycin treated WT and ‘del’ astrocytes compared to untreated; **b)** Western blot showing expression of membrane bound CHRNA7, CHRNA4 and cytoplasmic CHOP, p-PERK and PIK3R3 relative to GAPDH, internal control in untreated and tunicamycin treated WT and ‘del’ astrocytes. Note reduced expression of CHOP, p-PERK at protein level and *XBP1* splicing at RNA level in tunicamycin treated del’ astrocytes compared to treated WT astrocytes; **c)** volcano plot showing up (red upright triangles, n=236) and down (blue inverted triangles, n=255) regulated proteins (adjusted p-value≤0.05;±fold change) in tunicamycin treated ‘del’ astrocytes compared to treated WT astrocytes; **d)** GOCircle plot of differentially regulated proteins showing top 10 KEGG terms identified using DAVID functional annotation tool in i) untreated v/s tunicamycin treated WT astrocytes; and ii) tunicamycin treated WT v/s treated ‘del’ astrocytes. The inner ring represents a bar plot where height indicates the significance of the term (-log10 adjusted p-value), and color corresponds to the z-score. The outer ring displays scatterplots of the expression levels (logFC) for the genes in each term; and **e)** heat map constructed using selected proteins observed in GO terms at (d) above.

In transcriptome analysis, similar to untreated ‘del’ astrocytes (section 3.2 b), a significant increase in expression of various ion-channels, GPCRs, neurotransmitter receptors etc. via cAMP signaling (p-value≤0.05;±1 fold change) was seen in treated ‘del’ astrocytes compared to treated wild-type astrocytes (disease state), but the levels were only moderate (Supplementary Fig. 5a). Striking observation in the ‘del’ astrocytes under stress compared to disease state was reduced expression of known stress markers, namely CHOP (involved in apoptosis), phospho-PERK (inhibitor of translational initiation) at protein level; and lowered *XBP1* splicing (indicator of higher ER stress) at RNA level (Figs. 5a and b). This suggested reduced disease severity, likely modulated by either enhanced α4 nAChR surface expression or cAMP signaling in treated ‘del’ astrocytes under stress. To validate these findings, a comparative proteomic analysis was performed between wild-type and ‘del’ astrocytes treated with tunicamycin. A total of 236 upregulated and 255 downregulated proteins (adjusted p-value≤0.05;±1 fold change) (Fig. 5c) were identified in tunicamycin treated ‘del’ astrocytes. On enrichment analysis, many proteins which were downregulated in GO terms “SRP-dependent co-translational protein targeting to membrane”, “translational initiation” and “non-sense mediated decay” in disease state were now upregulated in tunicamycin treated ‘del’ astrocytes and *vice-versa* for KEGG terms “Amyotrophic lateral sclerosis”, “Parkinson disease”, “Huntington disease”, “Alzheimer’s disease”, “Spliceosome” (Fig. 5d and Supplementary Fig. 5b). These GO/KEGG terms included various small and large ribosomal subunits which were downregulated in disease state but upregulated in ‘del’ astrocytes under tunicamycin. On the other hand, several proteasome subunits; proteins CAPN2, CANX, CALR etc. involved in calcium homeostasis; and proteins TUBB and Rab subunits involved in neurodegenerative diseases, were reduced in tunicamycin treated ‘del’ astrocytes compared to disease state (Fig. 5e) with their log2FC values mentioned elsewhere (Supplementary Tables 5 and 6). Taken together, characterization of *RIC3* by gene based and comparative proteomic approaches suggest its likely protective/rescue role via nAChRs in the astrocyte disease model.

## 4. DISCUSSION

nAChRs are ligand-gated homo/hetero-pentameric ion channels involved in synaptic transmission and plasticity. Unlike other surface receptors, the assembly of these nAChRs in ER is a slow and inefficient process where each sub-unit undergoes post-translational modifications and adopts an appropriate functional confirmation ^27^. Dysregulation of nAChRs has been associated with a number of diseases including AD, PD, SZ, autism and attention deficit/hyperactivity disorder ^28^. Astrocytes, the most abundant non-neuronal cells involved in the pathogenesis of brain diseases also expresses these receptors ^13^. Furthermore, activation of nAChRs confers protection to astrocytes against oxidative stress, apoptosis and neuroinflammation ^11^. Selective molecular chaperones such as RIC-3 contribute to protein quality control of several of these nAChRs, but its role is rather poorly characterized in humans. This study aimed at characterization of *RIC3* using wild-type and Cas9 edited gene in human iPSC derived astrocytes (Fig. 1d) using targeted and hypothesis free analyses. Altered *RIC3* transcript ratio due to deletion induced splicing (Fig. 2a) and a gain of α7 nAChR surface expression in ‘del’ astrocytes (Fig. 2c) was noteworthy. Consequent reduced disease severity observed under tunicamycin induced ER stress in astrocyte disease model (Figs. 5b and d) suggests *RIC3* as a novel druggable target for nicotine related brain disorders. Salient findings of this study are discussed below.

### *RIC3* expression and implications for gene regulation

The first step to check for differences in expression, if any, of *RIC3* between wild-type and ‘del’ astrocytes revealed an unexpected extensive splicing alteration in ‘del’ astrocytes (Fig. 2a) compared to three annotated transcripts in wild-type astrocytes (Supplementary Fig. 2a). Of the six alternate transcripts in ‘del’ astrocytes (Fig. 2a), exon 2 skipping in a few may be explained by the 25bp deletion leading to loss of multiple ESE binding sites (Fig. 2c). Such splicing alterations are rare considering that large exonic deletions often result in frameshift and generation of a stop codon leading to mRNA degradation by non-sense mediated mRNA decay ^29^. However, newer studies show mRNA mis-regulation/splicing by disruption in ESE binding sites induced by CRISPR/Cas9 based mutagenesis ^30^; which lend support to splicing alteration observed in our study. Furthermore, the leaky nature of splicing witnessed by the mixture of transcripts including those with 25bp deletion; and an increased expression of shorter transcripts (Figs. 2a and b) reflects the complex RNA splicing regulatory mechanism induced by exonic deletions with therapeutic implications (discussed at the end). Deletion induced splicing resulting in i) presence of transcripts with 25bp deletion and exon 2 skipping predicted to result in frameshift and premature stop codon generation; and ii) presence as well as an increased expression of shorter *RIC3* transcripts (M5 and WT3) without second transmembrane and coiled-coil domains (Fig. 2a) leading to a gain of α7 nAChR surface expression in ‘del’ astrocytes was striking (Fig. 2c). Interestingly, the hypothesis free transcriptome analysis uncovered an increase in gene expression of various ion-channels, neurotransmitter receptors, GPCRs and cytokines (Figs. 3d and e). In addition, genes involved in cAMP, calcium/calmodulin-dependent kinase signaling and inflammatory responses controlled by transcription factors namely SRF, AP1, CEPB, NFKAPPB etc. were observed (Fig. 3c and Supplementary Fig. 3b).

*RIC3*, coding for an ER resident disordered chaperone protein belongs to a conserved gene family which is involved in folding and functional expression of nAChRs on surface ^31^. A conserved feature of RIC-3 across vertebrates and invertebrates is the presence of one or more coiled-coil domain which may be necessary for α7 nAChR maturation ^32,33^. Functional maturation of α7 nAChRs due to expression of RIC-3 isoforms without coiled-coil domain have been noted in both mammalian cell lines ^34^ and *Drosophila* ^35^. Furthermore, RIC-3 deletion analyses showed that ablation of the coiled-coil domain does not have any effect on expression of nAChRs ^36^. Alternative splicing in *RIC3* is seen across species, and in humans nine different protein coding transcripts including those without first and second transmembrane and coiled coil domains resulting from editing, alternative splicing and alternative promoters have been documented ^34^. These observations unequivocally support the functionality of annotated shorter *RIC3* transcript (WT3) encoding isoform without coiled-coil domain (c-termini) in the ‘del’ astrocytes. Both n-termini and c-termini of RIC-3 are predicted to be disordered region (Supplementary Fig. 6) and similar to other intrinsically disordered proteins, it can interact with or function as hubs in protein interaction networks regulating signaling pathways ^37^. This derive commendable support from another study wherein intracellular signaling proteins namely cAMP and inositol 1,4,5-trisphosphate receptor (IP3R) were identified in human α7 nAChR interactome, but only in the presence of RIC-3. The group also identified other proteins involved in biological processes such as post-translational processing of proteins and protein trafficking that affect nAChR surface expression ^38^. In addition, α7 nAChRs have been shown to co-localize with IP3Rs in PC12 cells ^39^ and the direct coupling of α7 nAChRs to G proteins can result in downstream calcium signaling response ^40^.

Therefore, we suggest a role of shorter RIC-3 isoforms as an adaptor resulting in i) the increased α7 nAChR expression through recruitment of proteins involved in receptor assembly, surface trafficking and modification; and ii) assembly/interactions of α7 nAChR with signaling proteins such as cAMP/GPCRs and their activation, in turn triggering alternate signaling cascades in ‘del’ astrocytes, seems likely. Importantly, the activation of cAMP and IP3R resulting in calcium mediated exocytosis of various cytokines and neurotransmitters can further trigger a series of intracellular processes resulting in signal amplification, by binding to their target receptors (neurotransmitters/GPCRs) on neighboring cells, as evident in ‘del’ astrocytes (Fig. 3b). However, due to unavailability of commercially available antibody against human RIC-3 that allow detection of multiple isoforms ^41^ and especially from c-termini, the functional role of shorter isoforms remain uncertain. Of note, another study highlights that the relevance of alterative splicing in *RIC3* is unclear and the shorter isoforms maybe non-functional ^42^. This may suggest that 25bp deletion induced mis-splicing results in loss-of-function and the increased α7 nAChR surface expression seen in ‘del’ astrocytes maybe due to upregulation of other chaperones, such as NACHO encoded by *TMEM35* ^43^. The marginal upregulation of *TMEM35* in ‘del’ astrocytes (log2FC=0.945) (Supplementary Table 3) may support this but warrants further analysis.

### *RIC3* and disease state

Functional relevance of the novel findings of gain of α7 nAChR expression and consequent activation of downstream signaling in ‘del’ astrocytes presented in ‘results’ (section 3.2 b) and discussed above were validated in the disease state. ER stress has been reported in various neurodegenerative diseases which occur due to accumulation of misfolded proteins or alterations in calcium homeostasis ^44^. Although such processes have been well described in various genetic and chemical induced disease models ^45,46^, the astrocyte-specific contribution to ER stress is poorly understood and our study findings seem to provide some leads.

#### Tunicamycin treated wild-type astrocytes

Disease relevant molecular perturbations observed in tunicamycin induced ER stress (Fig. 4) confirmed a successful generation of an astrocyte disease model for further analysis. Downregulation of various cell cycle and centromeric/mitotic spindle checkpoint genes (Figs. 4d and e), suggesting G2/M cell cycle arrest was most notable, providing newer insights into ER stress induced cell cycle arrest in astrocytes and subsequent apoptosis. In addition, induction of ER chaperones, upregulation of various proteasomal sub-units, genes/proteins involved in calcium homeostasis, translational attenuation, inhibition of non-sense mediated decay and reduced transport of receptors on membrane inferred from comparative transcriptome and proteomic analyses (Figs 4c; 5d and e), provide additional evidence of disease state. Interestingly, a significant upregulation of *RIC3* (Figs. 4e and f) and notable reduction in α7 nAChR levels but an increase in α4 nAChR subunits (Fig. 5b) which emerged as a salient feature encourages further discussion. These findings represent reported phenomena of i) higher chaperone levels under ER stress ^47^; and ii) aggregation of RIC-3 due to such an overexpression leading to reduced α7 nAChR surface expression ^48^. Higher *RIC3* levels in post-mortem brain samples of SZ and bipolar disorder ^49^ and loss of surface nAChRs in various neurodegenerative/neuropsychiatric disorders ^50^ also support our findings. In this context, it may be relevant to cite that mutations in *RIC3* were recently reported in PD. Limited functional analysis of these mutations indicated a likely role of *RIC3* in chaperone mediated receptor density alteration in PD ^51^ (OMIM#610509). However, the observed increased expression of α4 nAChR subunits in the tunicamycin treated wild-type astrocytes suggests a loss of inhibitory effects of RIC-3 under stress; and furthermore, that its interaction with α4 nAChR subtype maybe different, probably via different set of interacting proteins.

#### Tunicamycin treated ‘del’ astrocytes

A remarkable difference in our study was the much higher expression of α4 and not α7 nAChR subunits in tunicamycin treated ‘del’ astrocytes compared to disease state [(i) above] but with intact cAMP dependent signaling albeit minor quantitative reduction (Fig. 5b and Supplementary Fig. 5a). This may be interpreted to suggest contrasting effects of altered *RIC3* transcript ratio on different co-expressed nAChR subtypes under basal and stress condition. Alternately, shorter *RIC3* transcripts encoding isoforms with intact c-termini may result in interactions of α7 nAChRs with signaling proteins and their activation but not in their assembly/trafficking to surface under stress. On the other hand, the increased α4 nAChR subunit surface expression (Fig. 5b) and/or the intact signaling in ‘del’ astrocytes under stress i.e., increase in gene expression of various ion channels, neurotransmitter(s)/GPCRs (Supplementary Fig. 5a) may explain the reduced disease severity as evidenced by low levels of CHOP, p-PERK and *XBP1* splicing (Figs. 5a and b). Decrease in expression of proteasome subunits, proteins involved in protein processing in ER and various neurodegenerative diseases and upregulation of various ribosomal subunits identified in proteome analysis (Figs. 5d and e) additionally support decreased disease severity. Of note, under mild ER stress, unfolded protein response (UPR) has been reported to have a proadaptive role to restore protein homeostasis by inducing *XBP1* splicing (a potent transcriptional activator) and upregulation of UPR-targeted genes that not only increase the capacity of cells for protein folding, but also protein degradation and transport pathways. However, when the adaptive response fails, the sustained activation of PERK, an inhibitor of translation leads to upregulation of CHOP, a transcription factor implicated in the regulation of apoptosis ^52^. A similar upregulation was witnessed in disease state but lowered in ‘del’ astrocytes (Fig. 5b) suggesting a protective role of surface receptors/cAMP signaling under ER stress condition in astrocytes. Nicotine, a protective drug in PD has also been shown to upregulate α4β2 nAChRs ^53^, which may be conferring protection against cellular stress. Various drugs targeting GPCRs used to modulate canonical and noncanonical signaling pathways involved in neurodegenerative diseases ^54^ further support our hypothesis.

In summary, *RIC3*, a selective chaperone of nAChRs was characterized in astrocytes under basal and stress induced conditions utilizing a CRISPR/Cas9 mediated *RIC3* isogenic deletion iPSC line. The main findings included i) 25bp homozygous deletion in exon 2 resulting in splicing and altered *RIC3* transcript ratio; ii) gain of α7 expression and downstream signaling in ‘del’ astrocytes; and iii) splicing alteration having contrasting effects on co-expressed nAChRs with notable increase in α4 nAChR subunits under stress. In addition, a higher expression of shorter *RIC3* transcripts encoding isoforms with only c-termini may suggest its likely role as an adaptor with implications for peptidomimetic-based drug discovery in nicotine related brain disorders. Exonic deletion and not just splice junction mediated splicing alterations as captured in *RIC3* in this study may have additional implications. Akin to cellular rescue mechanism via deletion induced exon skipping as in the classical example of Duchenne muscular dystrophy ^55^, ASO based therapies may emerge for other disorders such as tauopathies.

## AUTHOR CONTRIBUTIONS

BKT designed the study and obtained research funding; NY performed the experiments; BKT and NY analyzed the results and wrote the first draft of the manuscript.

## ACKNOWLEDGEMENTS

We gratefully acknowledge Grant #BT/PR27457/MED/12/811/2018 to BKT and Senior Research Fellowship to NY from the Department of Biotechnology, Government of India, New Delhi; Junior and Senior Research Fellowship (#325541-2017-22) to NY from University Grants Commission (UGC), New Delhi; and Infrastructure support provided by UGC, New Delhi, through Special Assistance Programme and Department of Science and Technology, New Delhi, through FIST and DU-DST PURSE programmes to the Department of Genetics, University of Delhi South Campus. We thank Prof. Millet Treinin, Hebrew University, Israel, Dr. Antony Michealraj, University of Pittsburgh, USA and Dr. Sumedha Sudhaman, Vancouver, Canada for their invaluable comments and suggestions.

## Main Figure Legends

**Supplementary Fig. 1** Representative i) brightfield image of astrocytes showing star shaped structure (↑) at passage 2; and ii) phase contrast image of astrocytes showing mainly flattened polygonal to fusiform morphology at passage 4 (scale bar: 275μm).

**Supplementary Fig. 2 a)** Image showing nine annotated transcripts of human *RIC3* (green color) as retrieved from NCBI (https://www.ncbi.nlm.nih.gov/gene/79608?report=gene_table) with exons marked in red; **b)** Sanger sequencing confirmation of the gel eluted transcripts - WT2; WT3; and novel transcript M5 as described in Fig. 2a; and **c)** *in-silico* analysis using ESEfinder v3.0 and EX-SKIP showing loss of multiple exonic splicing enhancer binding sites due to 25bp deletion in exon 2 with a higher chance of exon skipping in ‘del’ astrocytes.

**Supplementary Fig. 3 a)** Enrichment analysis of differentially expressed genes in ‘del’ astrocytes compared to wild-type showing genes present in highly significant GO terms “plasma membrane”, “extracellular region”, “cytokine-mediated signaling pathway” and REACTOME terms “Neural system”, “Signaling by GPCR”, “Class A/1 (Rhodopsin-like receptors” and others (adjusted p-value≤0.05) identified using DAVID functional annotation tool. The color of the node and edge size correspond to the significance of the gene-set and number of genes overlapping between two connected gene-sets respectively; **b)** table showing number of genes regulated by transcription factors (adjusted p-value≤0.05) involved in different signaling pathways identified using UCSC_TFBS feature in DAVID.

**Supplementary Fig. 4** GOCircle plot of differentially regulated genes in tunicamycin treated WT astrocytes compared to untreated WT showing top 10 i) Biological processes and ii) Cellular component GO terms identified using DAVID functional annotation tool. The inner ring represents a bar plot where height indicates the significance of the term (-log10 adjusted p-value), and color corresponds to the z-score. The outer ring displays scatterplots of the expression levels (logFC) for the genes in each term.

**Supplementary Fig. 5 a)** Heat map constructed using selected significantly upregulated genes (p-value≤0.05;+1fold change) in tunicamycin treated ‘del’ astrocytes compared to tunicamycin treated wild-type (WT) astrocytes; and **b)** GOCircle plot of differentially regulated proteins showing top 10 Biological processes GO terms identified using DAVID functional annotation tool in i) untreated v/s tunicamycin treated WT astrocytes; and ii) tunicamycin treated WT v/s treated ‘del’ astrocytes. The inner ring represents a bar plot where height indicates the significance of the term (-log10 adjusted p-value), and color corresponds to the z-score. The outer ring displays scatterplots of the expression levels (logFC) for the genes in each term.

**Supplementary Fig. 6** Images showing **a)** Intrinsic disorder predisposition of human RIC-3 evaluated by PONDR®VSL2 and PONDR®VL-XT using on-line tool, PONDR (Predictor of Natural Disordered Regions; http://www.pondr.com/); and **b)** amino acids predicted to be disordered marked in blue box by DISOPRED3 (Disopred Prediction) using on-line tool, PSIPRED (http://bioinf.cs.ucl.ac.uk/psipred/). Note, the n-terminus, encompassing exon 1 (amino acids 1-42) and c-terminus having exon 6 (amino acids 223-368) regions are mostly disordered.

## REFERENCES

1. Schwappach, B. An overview of trafficking and assembly of neurotransmitter receptors and ion channels (Review). Mol. Membr. Biol. 25, 270–278 (2008).

2. Tao, Y.-X. & Conn, P. M. Chaperoning G protein-coupled receptors: from cell biology to therapeutics. Endocr. Rev. 35, 602–647 (2014).

3. Borroni, V. & Barrantes, F. J. Homomeric and Heteromeric α7 Nicotinic Acetylcholine Receptors in Health and Some Central Nervous System Diseases. Membranes (Basel). 11, (2021).

4. Albuquerque, E. X., Pereira, E. F. R., Alkondon, M. & Rogers, S. W. Mammalian nicotinic acetylcholine receptors: from structure to function. Physiol. Rev. 89, 73–120 (2009).

5. Buisson, B. & Bertrand, D. Nicotine addiction: the possible role of functional upregulation. Trends Pharmacol. Sci. 23, 130–136 (2002).

6. Caton, M., Ochoa, E. L. M. & Barrantes, F. J. The role of nicotinic cholinergic neurotransmission in delusional thinking. NPJ Schizophr. 6, 16 (2020).

7. Lombardo, S. & Maskos, U. Role of the nicotinic acetylcholine receptor in Alzheimer’s disease pathology and treatment. Neuropharmacology 96, 255–262 (2015).

8. Perez-Lloret, S. & Barrantes, F. J. Deficits in cholinergic neurotransmission and their clinical correlates in Parkinson’s disease. npj Park. Dis. 2, 16001 (2016).

9. Young, J. W. & Geyer, M. A. Evaluating the role of the alpha-7 nicotinic acetylcholine receptor in the pathophysiology and treatment of schizophrenia. Biochem. Pharmacol. 86, 1122–1132 (2013).

10. Jiwaji, Z. & Hardingham, G. E. Good, bad, and neglectful: Astrocyte changes in neurodegenerative disease. Free Radic. Biol. Med. 182, 93–99 (2022).

11. Sadigh-Eteghad, S., Majdi, A., Mahmoudi, J., Golzari, S. E. J. & Talebi, M. Astrocytic and microglial nicotinic acetylcholine receptors: an overlooked issue in Alzheimer’s disease. J. Neural Transm. 123, 1359–1367 (2016).

12. Liu, X. et al. Astrocytes in Neural Circuits: Key Factors in Synaptic Regulation and Potential Targets for Neurodevelopmental Disorders. Front. Mol. Neurosci. 14, 729273 (2021).

13. Wang, X., Lippi, G., Carlson, D. M. & Berg, D. K. Activation of α7-containing nicotinic receptors on astrocytes triggers AMPA receptor recruitment to glutamatergic synapses. J. Neurochem. 127, 632–643 (2013).

14. Yu, W.-F., Guan, Z.-Z., Bogdanovic, N. & Nordberg, A. High selective expression of alpha7 nicotinic receptors on astrocytes in the brains of patients with sporadic Alzheimer’s disease and patients carrying Swedish APP 670/671 mutation: a possible association with neuritic plaques. Exp. Neurol. 192, 215–225 (2005).

15. Hua, Y. et al. Activation of α7 Nicotinic Acetylcholine Receptor Protects Against 1-Methyl-4-Phenylpyridinium-Induced Astroglial Apoptosis. Front. Cell. Neurosci. 13, 507 (2019).

16. Liu, Y. et al. Activation of α7 nicotinic acetylcholine receptors protects astrocytes against oxidative stress-induced apoptosis: Implications for Parkinson’s disease. Neuropharmacology 91, 87–96 (2015).

17. Castillo, M. et al. Dual role of the RIC-3 protein in trafficking of serotonin and nicotinic acetylcholine receptors. J. Biol. Chem. 280, 27062–27068 (2005).

18. Lansdell, S. J. et al. RIC-3 enhances functional expression of multiple nicotinic acetylcholine receptor subtypes in mammalian cells. Mol. Pharmacol. 68, 1431–1438 (2005).

19. Millar, N. S. RIC-3: a nicotinic acetylcholine receptor chaperone. Br. J. Pharmacol. 153 Suppl, S177–83 (2008).

20. Silva, M. C. & Haggarty, S. J. Human pluripotent stem cell-derived models and drug screening in CNS precision medicine. Ann. N. Y. Acad. Sci. 1471, 18–56 (2020).

21. Kim, D., Paggi, J. M., Park, C., Bennett, C. & Salzberg, S. L. Graph-based genome alignment and genotyping with HISAT2 and HISAT-genotype. Nat. Biotechnol. 37, 907–915 (2019).

22. Love, M. I., Huber, W. & Anders, S. Moderated estimation of fold change and dispersion for RNA-seq data with DESeq2. Genome Biol. 15, 550 (2014).

23. Huang, D. W., Sherman, B. T. & Lempicki, R. A. Systematic and integrative analysis of large gene lists using DAVID bioinformatics resources. Nat. Protoc. 4, 44–57 (2009).

24. Walter, W., Sánchez-Cabo, F. & Ricote, M. GOplot: an R package for visually combining expression data with functional analysis. Bioinformatics 31, 2912–2914 (2015).

25. Smith, P. J. et al. An increased specificity score matrix for the prediction of SF2/ASF-specific exonic splicing enhancers. Hum. Mol. Genet. 15, 2490–2508 (2006).

26. Raponi, M. et al. Prediction of single-nucleotide substitutions that result in exon skipping: identification of a splicing silencer in BRCA1 exon 6. Hum. Mutat. 32, 436–444 (2011).

27. Green, W. N. & Millar, N. S. Ion-channel assembly. Trends Neurosci. 18, 280–287 (1995).

28. Hurst, R., Rollema, H. & Bertrand, D. Nicotinic acetylcholine receptors: from basic science to therapeutics. Pharmacol. Ther. 137, 22–54 (2013).

29. Doudna, J. A. & Charpentier, E. Genome editing. The new frontier of genome engineering with CRISPR-Cas9. Science 346, 1258096 (2014).

30. Tuladhar, R. et al. CRISPR-Cas9-based mutagenesis frequently provokes on-target mRNA misregulation. Nat. Commun. 10, 4056 (2019).

31. Treinin, M. RIC-3 and nicotinic acetylcholine receptors: Biogenesis, properties, and diversity. Biotechnol. J. 3, 1539–1547 (2008).

32. Biala, Y., Liewald, J. F., Ben-Ami, H. C., Gottschalk, A. & Treinin, M. The conserved RIC-3 coiled-coil domain mediates receptor-specific interactions with nicotinic acetylcholine receptors. Mol. Biol. Cell 20, 1419–1427 (2009).

33. Halevi, S. et al. Conservation within the RIC-3 gene family: Effectors of mammalian nicotinic acetylcholine receptor expression. J. Biol. Chem. 278, 34411–34417 (2003).

34. Seredenina, T., Ferraro, T., Terstappen, G. C., Caricasole, A. & Roncarati, R. Molecular cloning and characterization of a novel human variant of RIC-3, a putative chaperone of nicotinic acetylcholine receptors. Biosci. Rep. 28, 299–306 (2008).

35. Lansdell, S. J. et al. Host-cell specific effects of the nicotinic acetylcholine receptor chaperone RIC-3 revealed by a comparison of human and Drosophila RIC-3 homologues. J. Neurochem. 105, 1573–1581 (2008).

36. Ben-Ami, H. C. et al. RIC-3 affects properties and quantity of nicotinic acetylcholine receptors via a mechanism that does not require the coiled-coil domains. J. Biol. Chem. 280, 28053–28060 (2005).

37. Wright, P. E. & Dyson, H. J. Intrinsically disordered proteins in cellular signalling and regulation. Nat. Rev. Mol. Cell Biol. 16, 18–29 (2015).

38. Mulcahy, M. J., Blattman, S. B., Barrantes, F. J., Lukas, R. J. & Hawrot, E. Resistance to Inhibitors of Cholinesterase 3 (Ric-3) Expression Promotes Selective Protein Associations with the Human α7-Nicotinic Acetylcholine Receptor Interactome. PLoS One 10, e0134409 (2015).

39. Nordman, J. C. & Kabbani, N. Microtubule dynamics at the growth cone are mediated by α7 nicotinic receptor activation of a Gαq and IP3 receptor pathway. FASEB J. Off. Publ. Fed. Am. Soc. Exp. Biol. 28, 2995–3006 (2014).

40. King, J. R., Nordman, J. C., Bridges, S. P., Lin, M.-K. & Kabbani, N. Identification and Characterization of a G Protein-binding Cluster in α7 Nicotinic Acetylcholine Receptors. J. Biol. Chem. 290, 20060–20070 (2015).

41. Deshpande, A. et al. Why Does Knocking Out NACHO, But Not RIC3, Completely Block Expression of α7 Nicotinic Receptors in Mouse Brain? Biomolecules 10, (2020).

42. Halevi, S. et al. Conservation within the RIC-3 gene family. Effectors of mammalian nicotinic acetylcholine receptor expression. J. Biol. Chem. 278, 34411–34417 (2003).

43. Gu, S. et al. Brain α7 Nicotinic Acetylcholine Receptor Assembly Requires NACHO. Neuron 89, 948–955 (2016).

44. Lindholm, D., Wootz, H. & Korhonen, L. ER stress and neurodegenerative diseases. Cell Death Differ. 13, 385–392 (2006).

45. Cabral-Miranda, F. & Hetz, C. ER Stress and Neurodegenerative Disease: A Cause or Effect Relationship? Curr. Top. Microbiol. Immunol. 414, 131–157 (2018).

46. Oslowski, C. M. & Urano, F. Measuring ER stress and the unfolded protein response using mammalian tissue culture system. Methods Enzymol. 490, 71–92 (2011).

47. Ma, Y. & Hendershot, L. M. ER chaperone functions during normal and stress conditions. J. Chem. Neuroanat. 28, 51–65 (2004).

48. Shteingauz, A., Cohen, E., Biala, Y. & Treinin, M. The BTB-MATH protein BATH-42 interacts with RIC-3 to regulate maturation of nicotinic acetylcholine receptors. J. Cell Sci. 122, 807–812 (2009).

49. Severance, E. G. & Yolken, R. H. Lack of RIC-3 congruence with beta2 subunit-containing nicotinic acetylcholine receptors in bipolar disorder. Neuroscience 148, 454–460 (2007).

50. Dineley, K. T., Pandya, A. A. & Yakel, J. L. Nicotinic ACh receptors as therapeutic targets in CNS disorders. Trends Pharmacol. Sci. 36, 96–108 (2015).

51. Sudhaman, S. et al. Evidence of mutations in RIC3 acetylcholine receptor chaperone as a novel cause of autosomal-dominant Parkinson’s disease with non-motor phenotypes. J. Med. Genet. 53, 559–566 (2016).

52. Kadowaki, H. & Nishitoh, H. Signaling pathways from the endoplasmic reticulum and their roles in disease. Genes (Basel). 4, 306–333 (2013).

53. Srinivasan, R. et al. Nicotine up-regulates alpha4beta2 nicotinic receptors and ER exit sites via stoichiometry-dependent chaperoning. J. Gen. Physiol. 137, 59–79 (2011).

54. Azam, S. et al. G-Protein-Coupled Receptors in CNS: A Potential Therapeutic Target for Intervention in Neurodegenerative Disorders and Associated Cognitive Deficits. Cells 9, (2020).

55. Takeda, S., Clemens, P. R. & Hoffman, E. P. Exon-Skipping in Duchenne Muscular Dystrophy. J. Neuromuscul. Dis. 8, S343–S358 (2021).

