## Supplementary Figures for "Deletion induced splicing in *RIC3* drives nicotinic acetylcholine receptor regulation with implications for endoplasmic reticulum stress in human astrocytes"

**Supplementary Fig. 1**

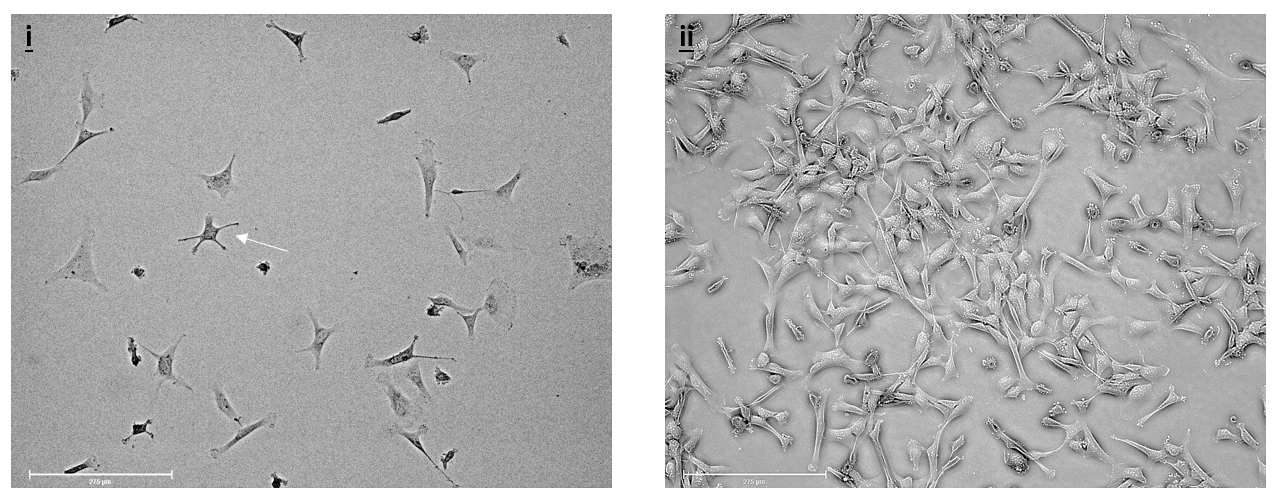

**Supplementary Fig. 2**

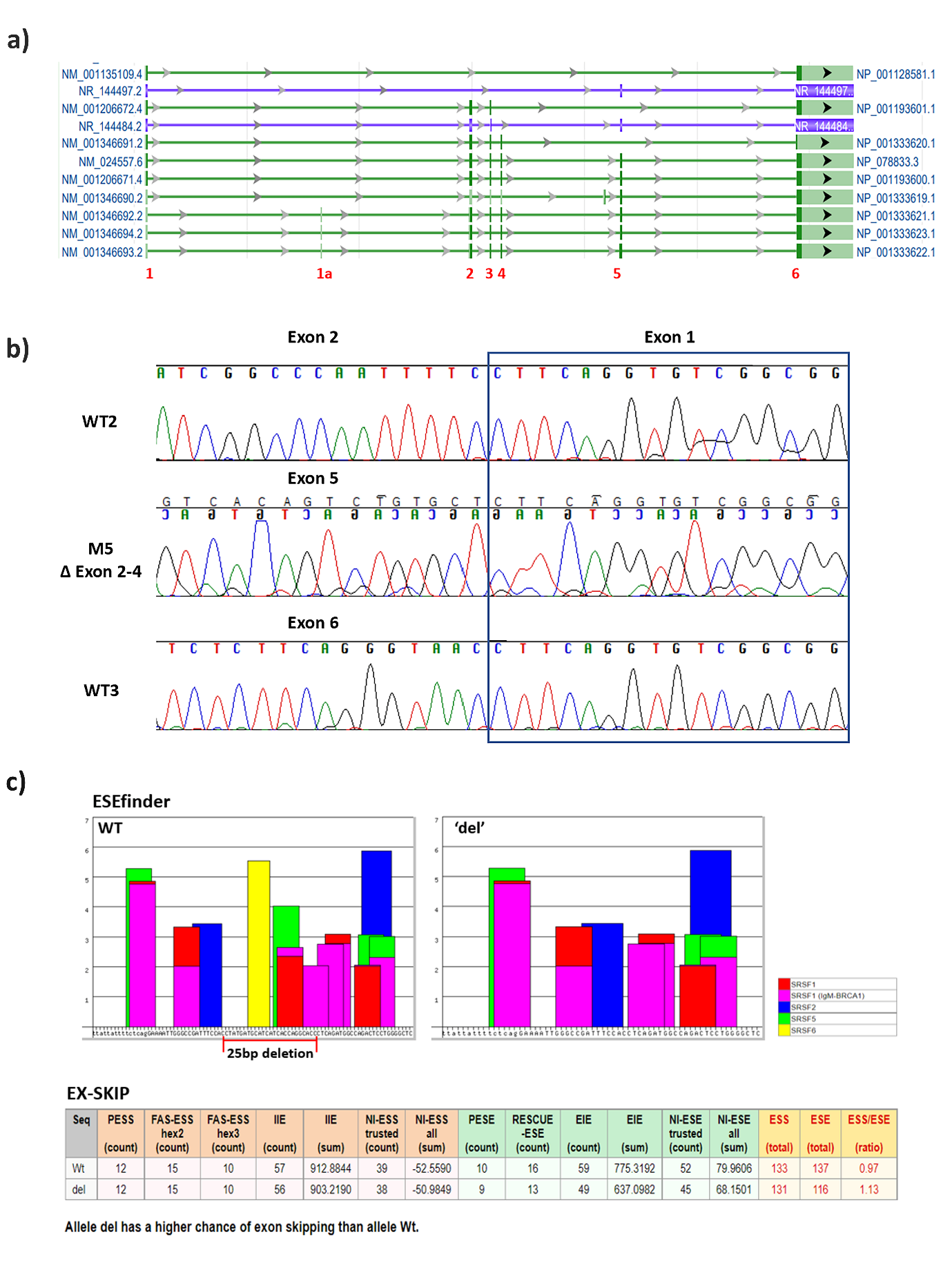

**Supplementary Fig. 3**

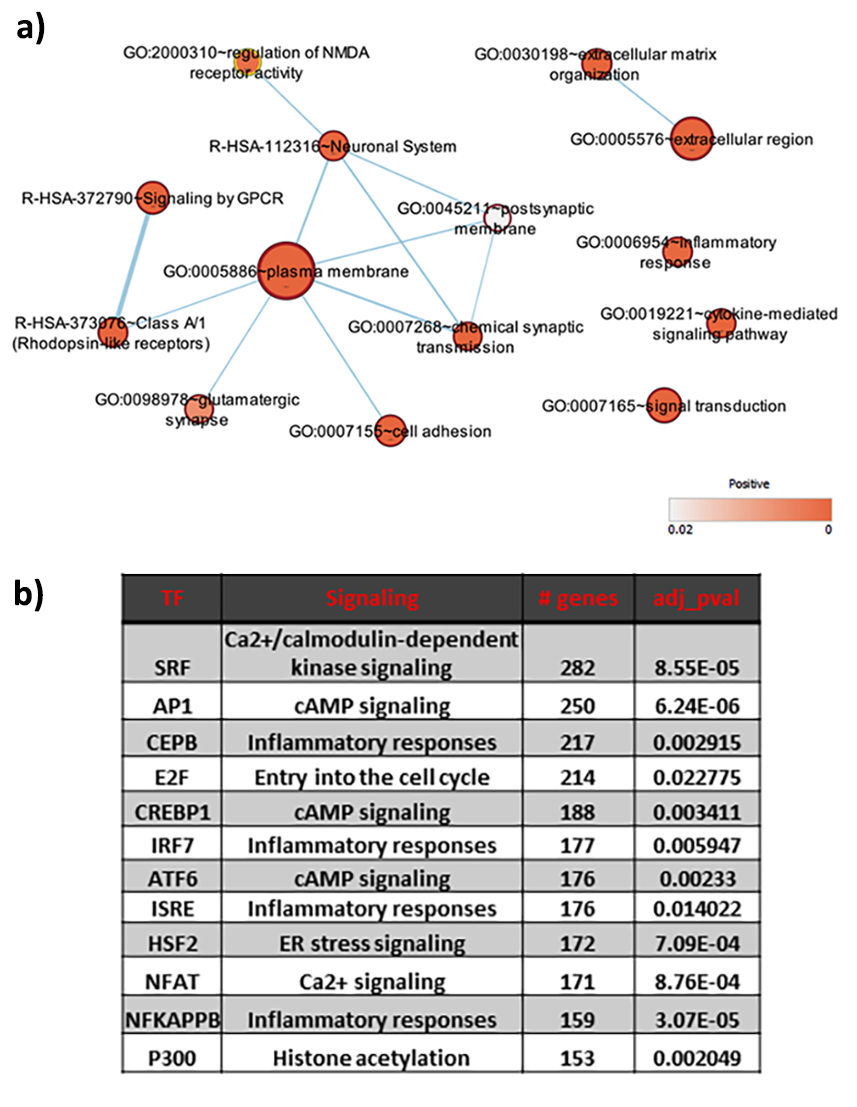

**Supplementary Fig. 4**

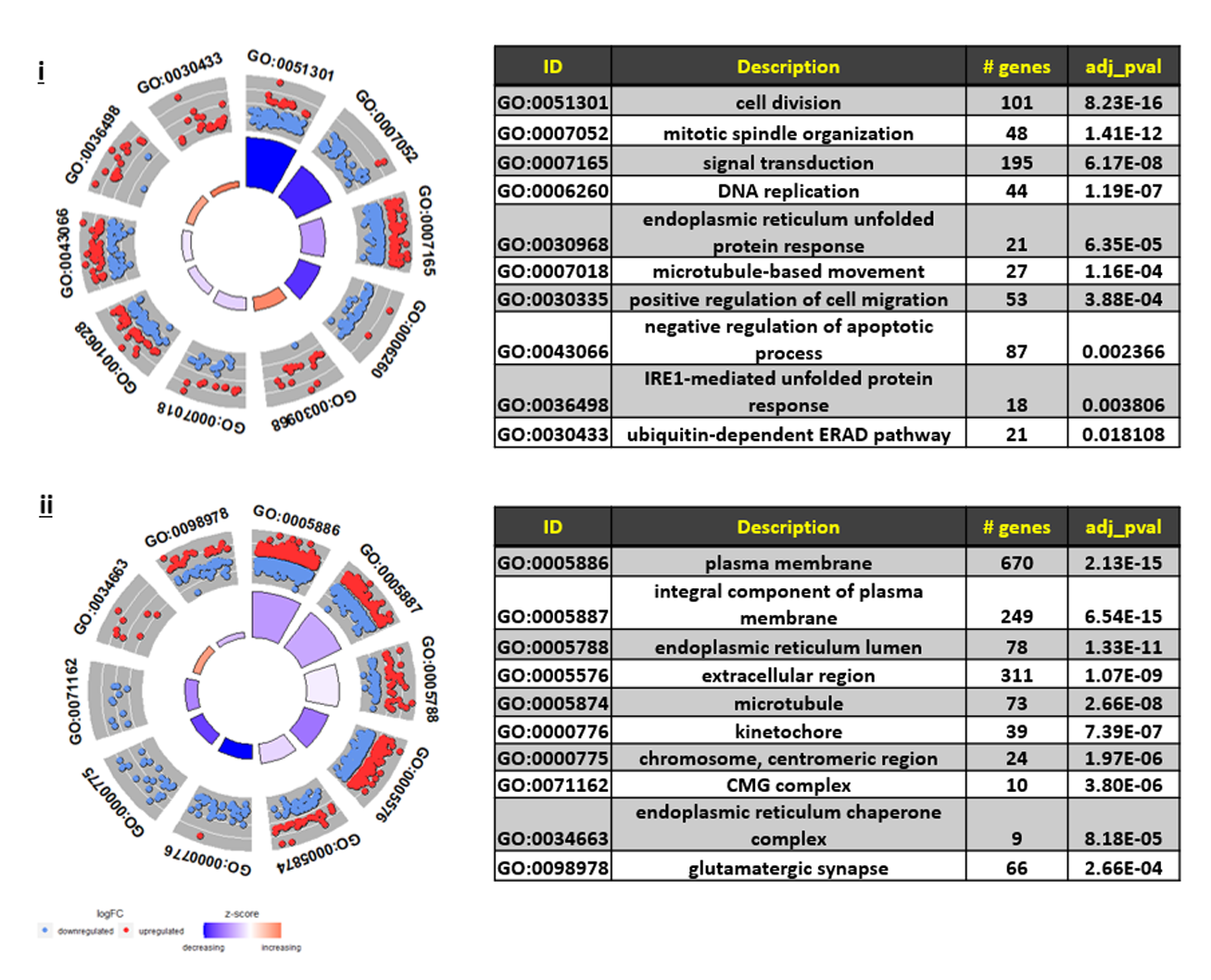

**Supplementary Fig. 5**

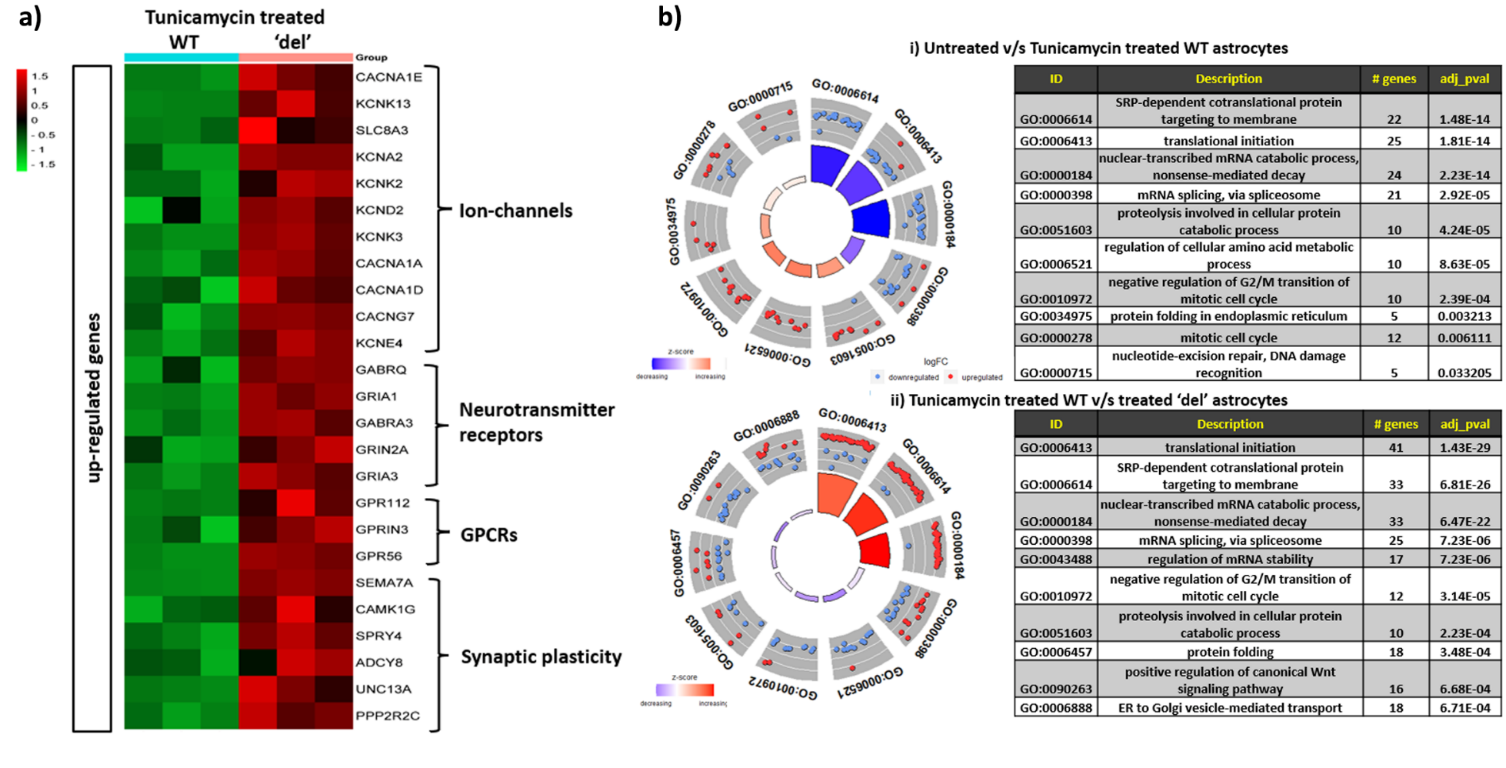

**Supplementary Fig. 6**

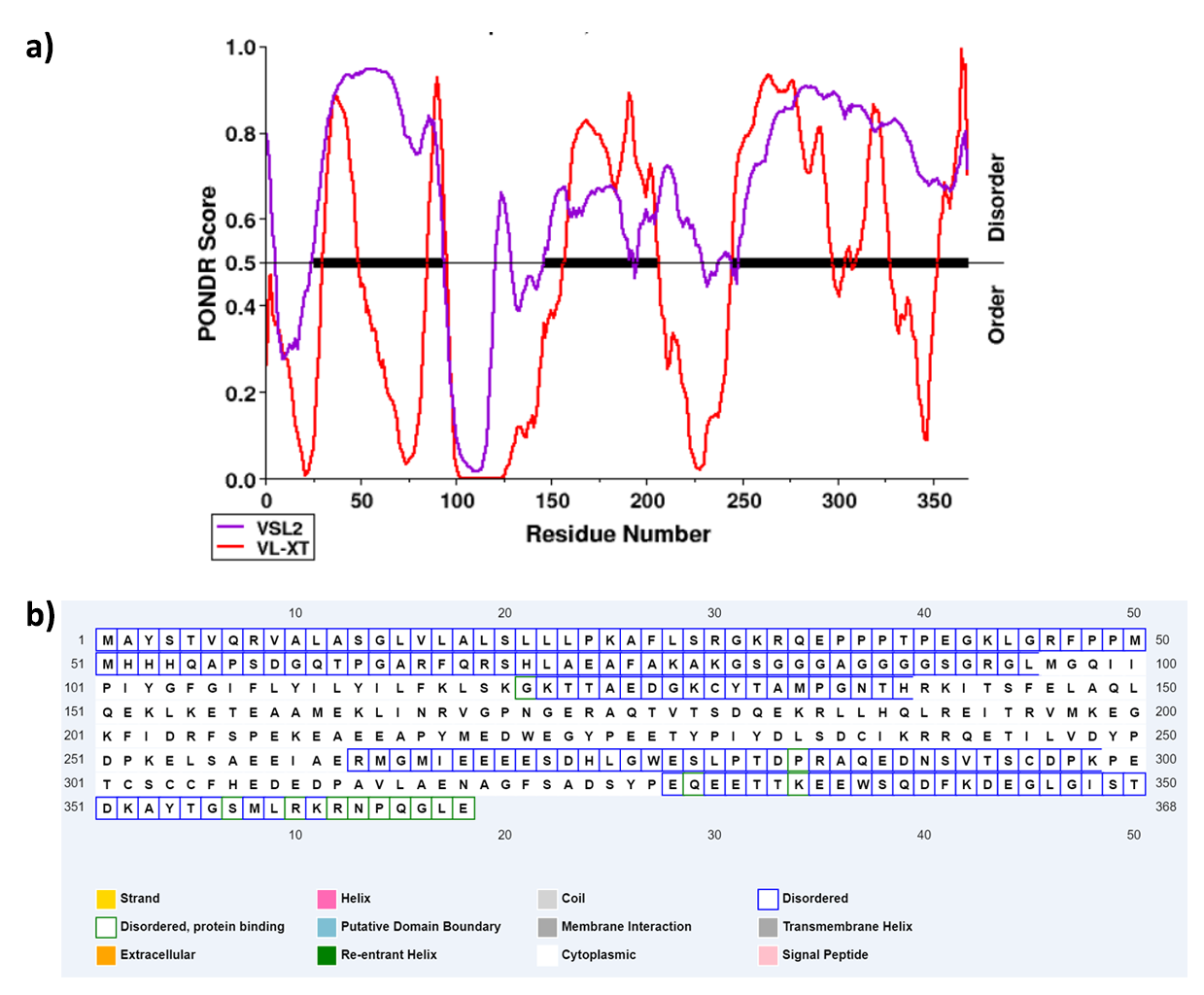
