## Supplementary Tables for "Deletion induced splicing in *RIC3* drives nicotinic acetylcholine receptor regulation with implications for endoplasmic reticulum stress in human astrocytes"

**Suppl. Table 1**: List of primer sequences used in the current study

| **Gene** | **Forward** | **Reverse** | **Application** |
| --- | --- | --- | --- |
| *RIC3* | GGGAAGAGAAAATGGGTTTGGT | CTCATGAAGAGGAGGCAGGATT | Sanger sequencing |
| *RIC3* | ATGGCGTACTCCACAGTGCAG | TCCTGTGGGTGTTTCCAGGCA | qPCR (E1-E3) |
| *RIC3* | ATGGCGTACTCCACAGTGCAG | CTGGCTTTGGATCACACGAGG | qPCR (E1-E6) |
| *RIC3* | CTTGTCCTGGCTCTGTCGCT | GGCTGCTTCTGTCTCCTTCAG | qPCR (E1-E4) |
| *XBP1S* | TTACGAGAGAAAACTCATGGC | GGGTCCAAGTTGTCCAGAATG | qPCR |
| *18S* | CGGCTACCACATCCAAGGAA | GCTGGAATTACCGCGGCT | RT-PCR |
| *CALR* | TTCGTTCTCAGTTCCGGCAAG | TGCTGAAAGGCTCGAAACTGG | RT-PCR |
| *CANX* | ACCACTGCTCCTCCTTCATCT | CGGTATCGTCTTTCTTGGCTT | RT-PCR |
| *CDC20* | AAAGCCAAGGAAGCCGCAG | CCAGGTTTGCTAGGAGTGGTC | RT-PCR |
| *CENPA* | CGCTTCCTCCCATCAACACAG | CCTGGGCTTGCCAATTGAAGT | RT-PCR |
| *CHOP* | CAGATGAAAATGGGGGTACCTA | GACTGGAATCTGGAGAGTGAGG | RT-PCR |
| *CHRM2* | CCAAGACCCCGTTTCTCCAAG | GCTCTCCTTCTCCTCTCCCTG | RT-PCR |
| *CHRNA1* | TCCTGGGCTCCGAACATGAG | TGTTTCAGACGCACATTGGTTG | RT-PCR |
| *GABBR2* | TCTGTCCATCCGTCACATCCA | ATTCACCGCATTGTCTGATGG | RT-PCR |
| *GABRQ* | GTCCCATCCCGAAATTCCACT | TCCAAAATTCGGTCTCAGGCG | RT-PCR |
| *GAPDH* | TGCACCACAACTGCTTAGC | GGCATGGACTGTGGTCATGAG | RT-PCR |
| *GRIA1* | AGGTGCCAATTTCCCCAACAA | GCTGTCGCTGATGTTCACAAT | RT-PCR |
| *GRIA3* | TTAGTCCTGGGGCTTTTGGGT | GCTTCTCGGTGGTGTTCTGG | RT-PCR |
| *GRIN2A* | CACACCTTCGTCCCCATCTTG | CCAGGGAGAAGACATGCCAGT | RT-PCR |
| *HSPA5* | GAGGTGGGCAAACAAAGACAT | CAGCAATAGTTCCAGCGTCTT | RT-PCR |
| *HSP40* | CATCAGAGCGCCAAATCAAGA | AGTAAAAGCACTGTGTCCAAGT | RT-PCR |
| *IL1B* | ACCTGAGCACCTTCTTTCCCT | GCGTGCAGTTCAGTGATCGTA | RT-PCR |
| *IL6R* | CAAAGGCTGTGCTCTTGGTGA | TTTTGCTGAACTTGCTCCCGA | RT-PCR |
| *MAPK13* | CATCAAGAAGCTGAGCCGACC | AGACATCCAGGAGCCCAATGA | RT-PCR |
| *MCM7* | TCTGTATGTGGACCTGGACGA | ACCACTTCCCTCTCCTTGTA | RT-PCR |
| *NESTIN* | GCGGGCTACTGAAAAGTTCCA | AGGCTGAGGGACATCTTGAGG | RT-PCR |
| *RIC3* | ATTACCAGTTTTGAGCTTGC | TCTCACCATTAGGTCCCACT | RT-PCR |
| *SNCA* | TGTAGGCTCCAAAACCAAGGA | CCTCCACTGTCTTCTGGGCTA | RT-PCR |
| *SOX1* | AACCCCAAGATGCACAACTCG | GCCAGCGAGTACTTGTCCTTC | RT-PCR |

**Supp. Table 2**: List and source of antibodies used for western blot and immunocytochemistry

| **Antibody** | **Host** | **Company #Cat No.** | **Dilution** | **Application** |
| --- | --- | --- | --- | --- |
| Anti-CHRNA4 | Mouse | Santa Cruz #sc-74519 | 1:1000 | Western blot |
| Anti-CHRNA7 | Goat | Everest #EB08853 | 1:1000 | Western blot |
| Anti-DDIT3/CHOP | Rabbit | ABclonal #A0221 | 1:1000 | Western blot |
| Anti-GAPDH | Rabbit | Abcam #ab9485 | 1:2500 | Western blot |
| Anti-GFAP | Rabbit | CST #12389T | 1:200 | Immunocytochemistry |
| Anti-NESTIN | Mouse | Invitrogen #A24345 | 1:50 | Immunocytochemistry |
| Anti-Phospho-PERK-T982 | Rabbit | ABclonal #AP0886 | 1:1000 | Western blot |
| Anti-PIK3R3/p55PIK | Rabbit | ABclonal #A3980 | 1:1000 | Western blot |
| Anti-RIC-3 | Rabbit | Invitrogen #PA5-111849 | 1:200 | Immunocytochemistry |
| Anti-SOX2 | Rabbit | Invitrogen #A24339 | 1:50 | Immunocytochemistry |
| HRP-conjugated anti-goat IgG | Donkey | Invitrogen #A15999 | 1:3000 | Western blot |
| HRP-conjugated anti-mouse IgG | Goat | Invitrogen #G21040 | 1:3000 | Western blot |
| HRP-conjugated anti-rabbit IgG | Goat | Abcam #ab6721 | 1:3000 | Western blot |
| Alexa Fluor 594 anti-rabbit | Donkey | Invitrogen #A24343 | 1:250 | Immunocytochemistry |
| Alexa Fluor 488 anti-mouse | Donkey | Invitrogen # A24350 | 1:250 | Immunocytochemistry |

**Suppl. Table 3**: List of differentially expressed genes in ‘del’ astrocytes compared to wild-type astrocytes under basal condition

| **Ion channels** | | | |
| --- | --- | --- | --- |
| **Ensembl ID** | **Gene** | **log2FC** | **padj** |
| ENSG00000100678 | *SLC8A3* | 2.307725994 | 1.36255E-24 |
| ENSG00000152315 | *KCNK13* | 2.27365571 | 0.048329748 |
| ENSG00000186510 | *CLCNKA* | 2.048859382 | 0.044813607 |
| ENSG00000177301 | *KCNA2* | 2.017639264 | 0.011383474 |
| ENSG00000141837 | *CACNA1A* | 1.806342669 | 3.4529E-43 |
| ENSG00000157551 | *KCNJ15* | 1.649373823 | 0.001626365 |
| ENSG00000137975 | *CLCA2* | 1.603838493 | 0.015547422 |
| ENSG00000148942 | *SLC5A12* | 1.581101334 | 1.22701E-05 |
| ENSG00000082482 | *KCNK2* | 1.524857009 | 5.44439E-11 |
| ENSG00000157388 | *CACNA1D* | 1.386562234 | 4.76626E-09 |
| ENSG00000116396 | *KCNC4* | 1.318996522 | 9.80248E-12 |
| ENSG00000189143 | *CLDN4* | 1.300812051 | 4.48137E-13 |
| ENSG00000171303 | *KCNK3* | 1.224418933 | 8.59846E-48 |
| ENSG00000184408 | *KCND2* | 1.16307194 | 0.000143271 |
| ENSG00000105605 | *CACNG7* | 1.15000729 | 8.82675E-07 |

| **Neurotransmitter genes** | | | |
| --- | --- | --- | --- |
| **Ensembl ID** | **Gene** | **log2FC** | **padj** |
| ENSG00000138435 | *CHRNA1* | 2.307919986 | 0.000239752 |
| ENSG00000183454 | *GRIN2A* | 1.774147391 | 1.20759E-05 |
| ENSG00000147402 | *GABRQ* | 1.462083476 | 2.13219E-09 |
| ENSG00000181072 | *CHRM2* | 1.435472489 | 0.008320958 |
| ENSG00000155511 | *GRIA1* | 1.051023508 | 3.66789E-07 |
| ENSG00000136928 | *GABBR2* | 1.015583484 | 1.6282E-11 |
| ENSG00000125675 | *GRIA3* | 1.000446947 | 6.24793E-39 |

| **G-protein coupled receptors** | | | |
| --- | --- | --- | --- |
| **Ensembl ID** | **Gene** | **log2FC** | **padj** |
| ENSG00000153292 | *GPR110* | 2.075038976 | 0.008029128 |
| ENSG00000069122 | *GPR116* | 1.8500089 | 9.90492E-05 |
| ENSG00000185477 | *GPRIN3* | 1.544518727 | 1.06246E-38 |
| ENSG00000183484 | *GPR132* | 1.420213459 | 1.09703E-05 |
| ENSG00000181773 | *GPR3* | 1.286516491 | 0.000157319 |
| ENSG00000205336 | *GPR56* | 1.134637574 | 2.50182E-56 |

| **Synaptic plasticity genes** | | | |
| --- | --- | --- | --- |
| **Ensembl ID** | **Gene** | **log2FC** | **padj** |
| ENSG00000163736 | *PPBP* | 4.026716801 | 0.019228212 |
| ENSG00000008118 | *CAMK1G* | 3.757891682 | 4.67E-27 |
| ENSG00000135447 | *PPP1R1A* | 3.049232412 | 0.010254264 |
| ENSG00000070808 | *CAMK2A* | 1.899352447 | 3.41E-40 |
| ENSG00000138623 | *SEMA7A* | 1.800972665 | 7.66E-167 |
| ENSG00000011347 | *SYT7* | 1.793590255 | 4.57E-05 |
| ENSG00000174600 | *CMKLR1* | 1.738062732 | 1.36E-05 |
| ENSG00000163888 | *CAMK2N2* | 1.485242686 | 0.008393493 |
| ENSG00000122025 | *FLT3* | 1.466023738 | 0.039003219 |
| ENSG00000156711 | *MAPK13* | 1.130009172 | 3.87E-06 |
| ENSG00000187678 | *SPRY4* | 1.093560305 | 1.81E-36 |
| ENSG00000130477 | *UNC13A* | 1.092903528 | 1.61E-29 |
| ENSG00000132639 | *SNAP25* | 1.087977308 | 4.89E-05 |
| ENSG00000107282 | *APBA1* | 1.074672547 | 5.32E-41 |
| ENSG00000078295 | *ADCY2* | 1.018803111 | 2.00E-10 |
| ENSG00000126950 | *TMEM35* | 0.94536859 | 0.029016363 |

| **Cytokines/receptors** | | | |
| --- | --- | --- | --- |
| **Ensembl ID** | **Gene** | **log2FC** | **padj** |
| ENSG00000162892 | *IL24* | 2.79843176 | 0.01162019 |
| ENSG00000174600 | *CMKLR1* | 1.738062732 | 1.36083E-05 |
| ENSG00000104951 | *IL4I1* | 1.245805187 | 2.95158E-06 |
| ENSG00000125538 | *IL1B* | 1.211806834 | 0.019545575 |
| ENSG00000160712 | *IL6R* | 1.046633382 | 0.004458639 |

| **Extracellular matrix genes** | | | |
| --- | --- | --- | --- |
| **Ensembl ID** | **Gene** | **log2FC** | **padj** |
| ENSG00000130702 | *LAMA5* | -1.003697462 | 0.000571611 |
| ENSG00000080573 | *COL5A3* | -1.066110901 | 9.12068E-23 |
| ENSG00000179403 | *VWA1* | -1.21778768 | 2.61378E-49 |
| ENSG00000166670 | *MMP10* | -1.346112376 | 0.025403358 |
| ENSG00000187955 | *COL14A1* | -1.352656137 | 1.42028E-12 |
| ENSG00000113361 | *CDH6* | -1.481694811 | 9.2884E-238 |
| ENSG00000182871 | *COL18A1* | -1.483014044 | 1.45785E-80 |
| ENSG00000112769 | *LAMA4* | -1.515601508 | 8.79363E-25 |
| ENSG00000196611 | *MMP1* | -1.683077417 | 0.013180983 |
| ENSG00000170801 | *HTRA3* | -2.151441609 | 2.03514E-58 |
| ENSG00000149968 | *MMP3* | -2.244673806 | 0.037570776 |
| ENSG00000189409 | *MMP23B* | -2.308562846 | 0.000804758 |
| ENSG00000137745 | *MMP13* | -2.32139139 | 7.80425E-18 |
| ENSG00000162692 | *VCAM1* | -2.351286492 | 8.75408E-16 |
| ENSG00000113209 | *PCDHB5* | -2.678008737 | 6.62664E-33 |
| ENSG00000171564 | *FGB* | -3.355944181 | 7.97217E-12 |
| ENSG00000150394 | *CDH8* | -6.118737621 | 5.66302E-32 |

**Suppl. Table 4**: List of differentially expressed genes in tunicamycin treated wild-type astrocytes compared to untreated wild-type astrocytes

| **Chaperones** | | | |
| --- | --- | --- | --- |
| **Ensembl ID** | **Gene** | **log2FC** | **padj** |
| ENSG00000166405 | *RIC3* | 2.540486478 | 9.47814E-16 |
| ENSG00000108176 | *DNAJC12* | 2.389696657 | 1.17783E-40 |
| ENSG00000173110 | *HSPA6* | 2.21133563 | 0.005345519 |
| ENSG00000113811 | *SELK* | 2.187979494 | 6.3956E-270 |
| ENSG00000162298 | *SYVN1* | 2.116267678 | 6.576E-106 |
| ENSG00000072849 | *DERL2* | 1.929925073 | 8.1378E-238 |
| ENSG00000116675 | *DNAJC6* | 1.683445999 | 2.90652E-70 |
| ENSG00000136770 | *DNAJC1* | 1.603422789 | 2.0785E-183 |
| ENSG00000135506 | *OS9* | 1.594903267 | 1.2715E-287 |
| ENSG00000077232 | *DNAJC10* | 1.452307285 | 6.1937E-257 |
| ENSG00000169087 | *HSPBAP1* | 1.355020218 | 1.26346E-29 |
| ENSG00000088298 | *EDEM2* | 1.327948048 | 1.03758E-98 |
| ENSG00000068912 | *ERLEC1* | 1.22249837 | 1.5414E-95 |
| ENSG00000185624 | *P4HB* | 1.189279077 | 5.1606E-131 |
| ENSG00000155304 | *HSPA13* | 1.153206203 | 1.1631E-115 |
| ENSG00000138942 | *RNF185* | 1.131603158 | 2.31392E-76 |
| ENSG00000197860 | *SGTB* | 1.06128061 | 5.51171E-59 |

| **Calcium homeostasis genes** | | | |
| --- | --- | --- | --- |
| **Ensembl ID** | **Gene** | **log2FC** | **padj** |
| ENSG00000110680 | *CALCA* | 5.292519768 | 3.71709E-09 |
| ENSG00000150995 | *ITPR1* | 1.58240972 | 2.53832E-71 |
| ENSG00000161921 | *CXCL16* | 1.524641183 | 8.45458E-06 |
| ENSG00000104805 | *NUCB1* | 1.514457825 | 6.3914E-141 |
| ENSG00000127022 | *CANX* | 1.472282024 | 1.9066E-231 |
| ENSG00000050393 | *MCUR1* | 1.182762241 | 4.01375E-85 |
| ENSG00000180879 | *SSR4* | 1.052789798 | 9.11454E-73 |
| ENSG00000074370 | *ATP2A3* | 1.040318607 | 9.39511E-06 |

| **Cell cycle genes** | | | |
| --- | --- | --- | --- |
| **Ensembl ID** | **Gene** | **log2FC** | **padj** |
| ENSG00000012048 | *BRCA1* | -1.037232219 | 8.80969E-23 |
| ENSG00000151849 | *CENPJ* | -1.055749621 | 5.04303E-26 |
| ENSG00000100297 | *MCM5* | -1.114278632 | 9.46895E-34 |
| ENSG00000170779 | *CDCA4* | -1.116981069 | 7.16818E-35 |
| ENSG00000120334 | *CENPL* | -1.125701915 | 1.76496E-16 |
| ENSG00000123374 | *CDK2* | -1.153179811 | 8.57089E-24 |
| ENSG00000163006 | *CCDC138* | -1.190049036 | 1.76191E-12 |
| ENSG00000097046 | *CDC7* | -1.197270638 | 3.76466E-17 |
| ENSG00000166508 | *MCM7* | -1.228497134 | 1.41181E-60 |
| ENSG00000125885 | *MCM8* | -1.271634753 | 9.70828E-41 |
| ENSG00000163814 | *CDCP1* | -1.274659987 | 1.5171E-106 |
| ENSG00000073111 | *MCM2* | -1.369054044 | 2.97978E-55 |
| ENSG00000076003 | *MCM6* | -1.437221333 | 6.51905E-90 |
| ENSG00000138092 | *CENPO* | -1.463815085 | 4.13059E-49 |
| ENSG00000123219 | *CENPK* | -1.570758436 | 1.19297E-31 |
| ENSG00000104738 | *MCM4* | -1.607961833 | 7.13692E-83 |
| ENSG00000139618 | *BRCA2* | -1.62672989 | 7.23147E-36 |
| ENSG00000166451 | *CENPN* | -1.749233042 | 2.69949E-49 |
| ENSG00000093009 | *CDC45* | -1.834912056 | 4.25776E-33 |
| ENSG00000144354 | *CDCA7* | -1.855995527 | 1.28668E-27 |
| ENSG00000153044 | *CENPH* | -1.868073719 | 1.67708E-32 |
| ENSG00000102384 | *CENPI* | -2.074371387 | 7.9595E-53 |
| ENSG00000151725 | *CENPU* | -2.192173461 | 5.13679E-52 |
| ENSG00000123080 | *CDKN2C* | -2.300169529 | 1.10187E-58 |
| ENSG00000203760 | *CENPW* | -2.659254788 | 1.87796E-74 |
| ENSG00000100526 | *CDKN3* | -2.784252046 | 2.79523E-68 |
| ENSG00000100162 | *CENPM* | -2.847319576 | 6.45226E-52 |
| ENSG00000158402 | *CDC25C* | -2.865363959 | 2.15104E-49 |
| ENSG00000111665 | *CDCA3* | -2.970279398 | 7.29107E-97 |
| ENSG00000146670 | *CDCA5* | -2.990533954 | 6.7687E-142 |
| ENSG00000138778 | *CENPE* | -3.019555228 | 6.23476E-79 |
| ENSG00000065328 | *MCM10* | -3.354676381 | 4.06545E-76 |
| ENSG00000117399 | *CDC20* | -3.43951141 | 9.5769E-85 |
| ENSG00000134690 | *CDCA8* | -3.441251566 | 1.4923E-150 |
| ENSG00000115163 | *CENPA* | -3.511158251 | 8.1893E-115 |
| ENSG00000170312 | *CDK1* | -3.579410747 | 2.5341E-239 |
| ENSG00000184661 | *CDCA2* | -3.639286062 | 2.727E-214 |

**Suppl. Table 5**: List of differentially expressed proteins in tunicamycin treated wild-type astrocytes compared to untreated wild-type astrocytes

| **Ribosomal subunits** | | | |
| --- | --- | --- | --- |
| **Uniprot ID** | **Protein** | **log2FC** | **padj** |
| P62273 | RPS29 | -2.35095 | 0.00475 |
| P62857 | RPS28 | -1.51984 | 0.005933 |
| P62899 | RPL31 | -1.42301 | 0.006035 |
| P83731 | RPL24 | -1.3471 | 0.013338 |
| P47914 | RPL29 | -1.34684 | 0.01326 |
| P46776 | RPL27A | -1.33951 | 0.01162 |
| P35268 | RPL22 | -1.32472 | 0.026998 |
| Q9Y3U8 | RPL36 | -1.31994 | 0.01569 |
| P62851 | RPS25 | -1.29143 | 0.013338 |
| P62266 | RPS23 | -1.26583 | 0.023576 |
| P61254 | RPL26 | -1.21822 | 0.013006 |

| **Proteasome subunits** | | | |
| --- | --- | --- | --- |
| **Uniprot ID** | **Protein** | **log2FC** | **padj** |
| Q9UNM6 | PSMD13 | 1.68481 | 0.044962 |
| O75832 | PSMD10 | 1.639196 | 0.009426 |
| P25789 | PSMA4 | 1.452012 | 0.007193 |
| P20618 | PSMB1 | 1.399922 | 0.023043 |
| P25787 | PSMA2 | 1.368497 | 0.023064 |
| P25788 | PSMA3 | 1.208344 | 0.014982 |
| Q99436 | PSMB7 | 1.195137 | 0.026487 |
| O00231 | PSMD11 | 1.192862 | 0.031301 |
| O43242 | PSMD3 | 1.005893 | 0.032531 |

| **Calcium homeostasis proteins** | | | |
| --- | --- | --- | --- |
| **Uniprot ID** | **Protein** | **log2FC** | **padj** |
| P16615 | ATP2A2 | 2.351817 | 0.005537 |
| P17655 | CAPN2 | 1.886522 | 0.009475 |
| P27824 | CANX | 1.555117 | 0.00475 |
| P27797 | CALR | 1.106458 | 0.031063 |

| **Tubulin subunits** | | | |
| --- | --- | --- | --- |
| **Uniprot ID** | **Protein** | **log2FC** | **padj** |
| Q9BVA1 | TUBB2B | 1.955878 | 0.00475 |
| Q9BUF5 | TUBB6 | 1.319131 | 0.005552 |
| P07437 | TUBB | 1.221814 | 0.007193 |
| Q13509 | TUBB3 | 1.120793 | 0.008452 |
| P04350 | TUBB4A | 1.08069 | 0.019762 |

| **Rab subunits** | | | |
| --- | --- | --- | --- |
| **Uniprot ID** | **Protein** | **log2FC** | **padj** |
| Q15907 | RAB11B | 1.819156 | 0.00734 |
| P20339 | RAB5A | 1.784528 | 0.004375 |

**Suppl. Table 6**: List of differentially expressed proteins in tunicamycin treated ‘del’ astrocytes compared to treated wild-type astrocytes

| **Ribosomal subunits** | | | |
| --- | --- | --- | --- |
| **Uniprot ID** | **Protein** | **log2FC** | **padj** |
| P62857 | RPS28 | 1.883172 | 0.005366 |
| P49207 | RPL34 | 1.829582 | 0.00354 |
| P62899 | RPL31 | 1.711144 | 0.00354 |
| P42766 | RPL35 | 1.550387 | 0.00789 |
| P62273 | RPS29 | 1.541193 | 0.008671 |
| P47914 | RPL29 | 1.532405 | 0.010552 |
| P46778 | RPL21 | 1.496812 | 0.012233 |
| P62753 | RPS6 | 1.484212 | 0.003813 |
| P83881 | RPL36A | 1.476419 | 0.013031 |
| Q02543 | RPL18A | 1.463555 | 0.021618 |
| P39019 | RPS19 | 1.453891 | 0.004056 |
| P61254 | RPL26 | 1.433906 | 0.007077 |
| P62917 | RPL8 | 1.433751 | 0.009114 |

| **Proteasome subunits** | | | |
| --- | --- | --- | --- |
| **Uniprot ID** | **Protein** | **log2FC** | **padj** |
| P28066 | PSMA5 | -1.96921 | 0.012233 |
| P25787 | PSMA2 | -1.88748 | 0.016161 |
| O43242 | PSMD3 | -1.84363 | 0.008005 |
| P25788 | PSMA3 | -1.80372 | 0.00659 |
| Q9UL46 | PSME2 | -1.52274 | 0.004011 |
| Q99436 | PSMB7 | -1.49923 | 0.019614 |
| O75832 | PSMD10 | -1.32978 | 0.011547 |
| P25789 | PSMA4 | -1.30578 | 0.016901 |
| Q06323 | PSME1 | -1.24606 | 0.025944 |
| P62333 | PSMC6 | -1.23349 | 0.004056 |

| **Calcium homeostasis proteins** | | | |
| --- | --- | --- | --- |
| **Uniprot ID** | **Protein** | **log2FC** | **padj** |
| P17655 | CAPN2 | -4.0543 | 0.008376 |
| P16615 | ATP2A2 | -3.53567 | 0.003201 |
| P27824 | CANX | -2.66781 | 0.00354 |
| P27797 | CALR | -1.3415 | 0.011058 |

| **Tubulin subunits** | | | |
| --- | --- | --- | --- |
| **Uniprot ID** | **Protein** | **log2FC** | **padj** |
| Q9BVA1 | TUBB2B | -1.81675 | 0.00354 |
| Q13509 | TUBB3 | -1.31337 | 0.006218 |
| P04350 | TUBB4A | -1.21831 | 0.013871 |
| Q9BUF5 | TUBB6 | -1.21625 | 0.011855 |
| P07437 | TUBB | -1.10387 | 0.007154 |

| **Rab subunits** | | | |
| --- | --- | --- | --- |
| **Uniprot ID** | **Protein** | **log2FC** | **padj** |
| P61026 | RAB10 | -3.02453 | 0.00354 |
| Q15907 | RAB11B | -2.95843 | 0.003618 |
| P51149 | RAB7A | -2.01067 | 0.022999 |
| P62820 | RAB1A | -1.44717 | 0.006668 |
